# Age-associated downregulation of glutamate and GABA neurotransmission-related gene expression in the rostral ventrolateral medulla of male Fischer 344 rats

**DOI:** 10.1101/2022.12.14.520496

**Authors:** Sivasai Balivada, Geronimo P. Tapia, Hitesh N. Pawar, Arshad M. Khan, Michael J. Kenney

## Abstract

The rostral ventrolateral medulla (RVLM), a part of the medullary reticular formation, plays a major role in several physiological responses, including cardiovascular and sympathetic nervous system functions. Although aging causes disturbances in the responses of these physiological systems, RVLM involvement in these age-related changes is not clear. Previous work using high-throughput gene expression analysis of the RVLM in aged animals suggested that chemical neurotransmission-related genes might be downregulated with advancing age. Since RVLM function involves a balance of signals from inhibitory and excitatory inputs, which is largely mediated by gamma-aminobutyric acid (GABA) and excitatory amino acid (EAA) neurotransmission, we hypothesized that aging is associated with altered excitatory and/or inhibitory neurotransmission-related gene expression in the RVLM. To test this hypothesis, we micropunched an RVLM-containing area from young (3–5 months), middle-aged (12–14 months), and aged (22– 26 months) Fischer 344 male rats. RNA purified from these micropunches was analyzed using GABA and Glutamate RT^2^ Profiler PCR arrays (n= 8–10). Each profiler array has primers for 84 GABA and glutamate neurotransmission related genes. In addition, the expression of selected genes was validated at the RNA level using TaqMan^®^ based-qPCR and at the protein level using western blotting. All the genes that displayed significant differential expression (1.5-fold, *p* < .05, FDR < .05) were identified to be downregulated in the RVLM of aged and middle-aged rats compared to young rats. This downregulation did not appear to be a result of RVLM tissue sampling differences among the age groups, since a separate validation of our sampling method, which involved careful mapping of micropunched regions to a standardized brain atlas, revealed no spatial differences in sampled sites among age groups. Among the downregulated genes, the percentage of glutamate neurotransmission-related genes was higher than GABA neurotransmission-related genes. The Solute carrier family 1 member 6 (*Slc1a6*) gene showed the highest fold downregulation at the RNA level in the RVLM of aged compared to young rats, and its protein product, Excitatory amino acid transporter 4 (EAAT4), showed a downregulatory trend in the RVLM of aged and middle-aged rats. These results suggest that molecular constituents of both GABA and glutamate neurotransmission might be altered in the RVLM of aged and middle-aged rats, and the changes in glutamate neurotransmission might be more prominent. Investigating age-associated anatomical and functional changes in RVLM GABA and glutamate neurotransmission might provide a foundation for understanding the effects of aging on physiological function.

## Introduction

The rostral ventrolateral medulla (RVLM) is a lower brainstem area that contains diverse neural substrates involved in the regulation of multiple physiological systems, including the respiratory, cardiovascular (CV) and the sympathetic nervous system (SNS) (Blessing, 1997; Schreihofer and Sved, 2011). A foundational tenet of RVLM function is that it involves a balance of inhibitory and excitatory input signals, mediated to a large extent by gamma-amino butyric acid (GABA) and excitatory amino acid (EAA) neurotransmission, with a particularly striking effect of this balance on CV and SNS regulation (Ito and Sved, 1997; Kenney, 2014; Kenney et al., 2011; Ross et al., 1984; Sun and Guyenet, 1985). For example, microinjections of glutamate receptor agonists into the RVLM increase arterial blood pressure and sympathetic nerve activity in conscious (Sakima et al., 2000) and anesthetized rats (Kenney, 2014; Kenney et al., 2013; Kenney et al., 2011; Ross et al., 1984). Similarly, microinjections of GABA receptor antagonists in the RVLM increase arterial blood pressure and sympathetic nerve activity (Kenney, 2014; Kenney et al., 2013; Sun and Guyenet, 1985), while administration of GABA receptor agonists decrease arterial blood pressure and sympathetic nerve activity (Kenney et al., 2011; Ross et al., 1984). Moreover, alterations in RVLM GABAergic and/or glutamatergic neurotransmission have been observed in conditions known to be associated with altered CV and SNS regulation, such as chronic heart failure (Wang et al., 2009), hypertension (Smith and Barron, 1990), obesity (Huber and Schreihofer, 2011; Ito et al., 2001), sedentary behavior (Mueller, 2007; Mueller and Mischel, 2012), and advanced age (Kenney, 2014).

Advancing age in the absence of clinical disease is associated with numerous changes in SNS regulation (Kenney, 2010; Seals and Esler, 2000), including: alterations in the sympathetic neural regulation of arterial blood pressure, changes in the background level of sympathetic nerve outflow, and modifications in the responsiveness of the SNS to various stressors (Floras, 2021; Kenney, 2010; McCarty, 1986; Seals and Esler, 2000). A combination of factors provides a strong rationale for seeking to understand central neural mechanisms mediating age-related changes in SNS regulation. These include the prominent increase in the aging population, age-related changes in sympathetic nerve outflow and responsivity, the critical relationship between the SNS and physiological function, and the contribution of SNS dysregulation to the development of chronic disease (Kenney, 2010). With these issues in mind, we have completed a series of studies evaluating whether aging alters the molecular and cellular substrate of the RVLM (Balivada et al., 2017, 2019, 2021; Pawar et al., 2017, 2018).

Previously, we analyzed the basal RNA expression of select GABA and glutamate ionotropic receptor subunits in the RVLM-containing area of young and aged Fischer 344 (F344) rats using quantitative reverse transcription polymerase chain reaction (qRT-PCR)-based methods (Pawar et al., 2017, 2018). No significant age-related differences in the RVLM expression of these receptor subunit genes were found. In a subsequent study, Balivada et al., (2017) analyzed global gene expression patterns in the RVLM of young, middle-aged, and aged F344 rats at the RNA level using gene ontology and pathway analyses. The expression patterns revealed an age-associated decrease in chemical neurotransmission-related genes in the RVLM of aged rats at the RNA level (Balivada et al., 2017). Specifically, genes related to the neurotransmitter metabolism/release cycle and synaptic plasticity were downregulated in the RVLM of aged rats (Balivada et al., 2017), modifications that may provide the molecular backdrop for understanding age-dependent changes in RVLM function. However, these two studies were limited in at least three ways. First, Pawar et al. (2017 and 2018) used a two-fold change in gene expression as the threshold for detecting differences between aged vs. young RVLM and it is possible that there might be gene expression differences that are less than two-fold in magnitude. In fact, several studies have used the combination of less than two-fold difference with statistical significance as a framework to establish changes in gene expression (Peart et al., 2005; Raouf et al., 2008). Second, a potential effect of aging on GABA and glutamate neurotransmission-related gene expression in the RVLM of middle-aged F344 rats was not explored. Third, results from the global gene expression analysis did not address whether age-associated changes in neurotransmission-related genes were related to RVLM GABAergic or glutamatergic systems, or both. Importantly, the effect of age on specific aspects of GABAergic and glutamatergic neurotransmission in the RVLM, such as metabotropic receptors, transporters, trafficking proteins, and downstream signaling proteins remain unknown.

In the present study, we characterized the transcriptomic profile of RVLM GABA and glutamate neurotransmission-related genes to test the hypothesis that aging is associated with altered excitatory and/or inhibitory neurotransmission-related gene expression in the RVLM. The RVLM-containing lower brainstem area was micropunched from young, middle-aged, and aged F344 rats and purified RNA was analyzed using RT^2^ profiler arrays which contained genes related to GABAergic and glutamatergic-neurotransmission, including: ionotropic and metabotropic neurotransmitter receptors, membrane and vesicular transporters, trafficking proteins, proteins involved in GABA and glutamate metabolism, and downstream signaling molecules. Alterations in the expression of select genes were validated at the mRNA level using TaqMan^®^ based-qPCR and at the protein level using immunoblotting. Moreover, our sampling methodology was also validated using an atlas-based mapping approach (Khan et al., 2018). We observed a decrease in select GABA and glutamate neurotransmission-related gene expression in the RVLM of middle-aged and aged rats compared to young rats, with a particular age-related influence on genes involved in glutamate neurotransmission.

## Methods

### Animals and tissue collection

Male Fischer 344 (F344) rats obtained from Charles River Laboratories (contracted with the National Institute on Aging) were used, as they have been extensively employed for investigating aging effects on animal health (Nadon, 2006; Mitchell et al., 2015). All procedures involving F344 rats in this study were approved by the Institutional Animal Care and Use Committee. Rats 3–5 months old (304 ± 7 g), 12–14 months old (425 ± 8 g), and 22–26 months old (432 ± 11 g) were categorized as young, middle-aged, and aged, respectively. Rats were housed in a conventional housing facility in a temperature-controlled room at 24°C and on a 12:12 h light-dark cycle. Lights were turned on at 7:00 AM and turned off at 7:00 PM. The body weight of the young rats was significantly different from that of the middle-aged and aged rat groups (ANOVA, Tukey HSD *p* < 0.05), and there was no difference in body weight between middle-aged and aged rats. The body weights of the aged rats were measured every week and the animals that showed progressive declines in their body weight were not included in the study. On the day of the tissue harvest, rats were deeply anesthetized with 5% isoflurane (IsoSol, Vedco, Inc., Saint Joseph, MO, USA) in an anesthetic induction chamber for a minimum of 5 min until the absence of reflex to mechanical stimulation of the hindlimb. Deeply anesthetized rats were euthanized by decapitation with a guillotine. Brains and liver samples were collected immediately, snap-frozen with liquid nitrogen in a polypropylene vial, and stored at −80°C until further use. Tissue harvests were performed between 9:00 AM and 5:00 PM.

### Micropunching procedure of the RVLM area

The RVLM-containing area was micropunched as described by Balivada et al., (2017) with some modifications to the procedure as noted here. The stereotaxic coordinate system of the rat brain atlas by Paxinos & Watson (*PW7*) (Paxinos and Watson, 2014), along with its photographic documentation of histologically stained hindbrain sections, served as the set of references to locate and identify the hindbrain region containing the RVLM. For sample collection purposes, we defined that the ventrolateral 0.79 mm^2^ area (diameter-1 mm) beside the spinal trigeminal tract (sp5) and the olivocerebellar tract (oc) on Figs. 133 (inferred Bregma distance of −12.00 mm) through 137 (inferred Bregma distance of −12.48 mm) of the *PW7* atlas contains the RVLM. Based on this definition, we used inferred stereotaxic space (Paxinos and Watson, 2014; section 4.6.2.1.1 in Khan (2013) for more on the concept) to perform calibrated sectioning. To collect hindbrain sections containing the RVLM area, coronal sections were made through the rostrocaudal extent of the brain using a cryostat (CM3050S, Leica Biosystems, IL, USA). After adjusting the horizontal orientation (**Figure 1a**) we identified lobule 2 of the cerebellar vermis (abbreviated ’2Cb’; Figure 103 of *PW7* (the rostral fiducial that was used to mark the beginning of calibrated sectioning; **Figure 1b**). Ninety 40 μm-thick serial sections (∼3.6 mm) were obtained from the point of the lobule’s appearance in the tissue and then discarded. From the remaining tissue block-face, two serial 200 μm-thick coronal sections were collected (**Figure 1c**) (beginning at an inferred distance from Bregma of −12.0 mm, or approximately at Figure 133, and extending to −12.4 mm of Bregma, or approximately at the level shown in Figure 136 of *PW7*). In these sections, the RVLM-containing area was micropunched bilaterally using a Harris micropunch (inner diameter = 1 mm) on the cryostat blade holder (cooled to −20°C) following a modification of the Palkovits micropunch technique (Palkovits, 1985). All the four tissue punches from each brain were pooled (tissue volume of ∼0.8 mm^3^) into an RNase-free centrifuge tube and stored at −80°C until future use. After processing each brain, the micropunch tip was cleaned with 100% ethanol followed by application of RNase Zap (Thermo Fisher Scientific, Waltham, MA, USA) to prevent cross-contamination. The cryostat blade holder was cleaned with 100% ethanol prior to cryosectioning.

**Figure 1:**
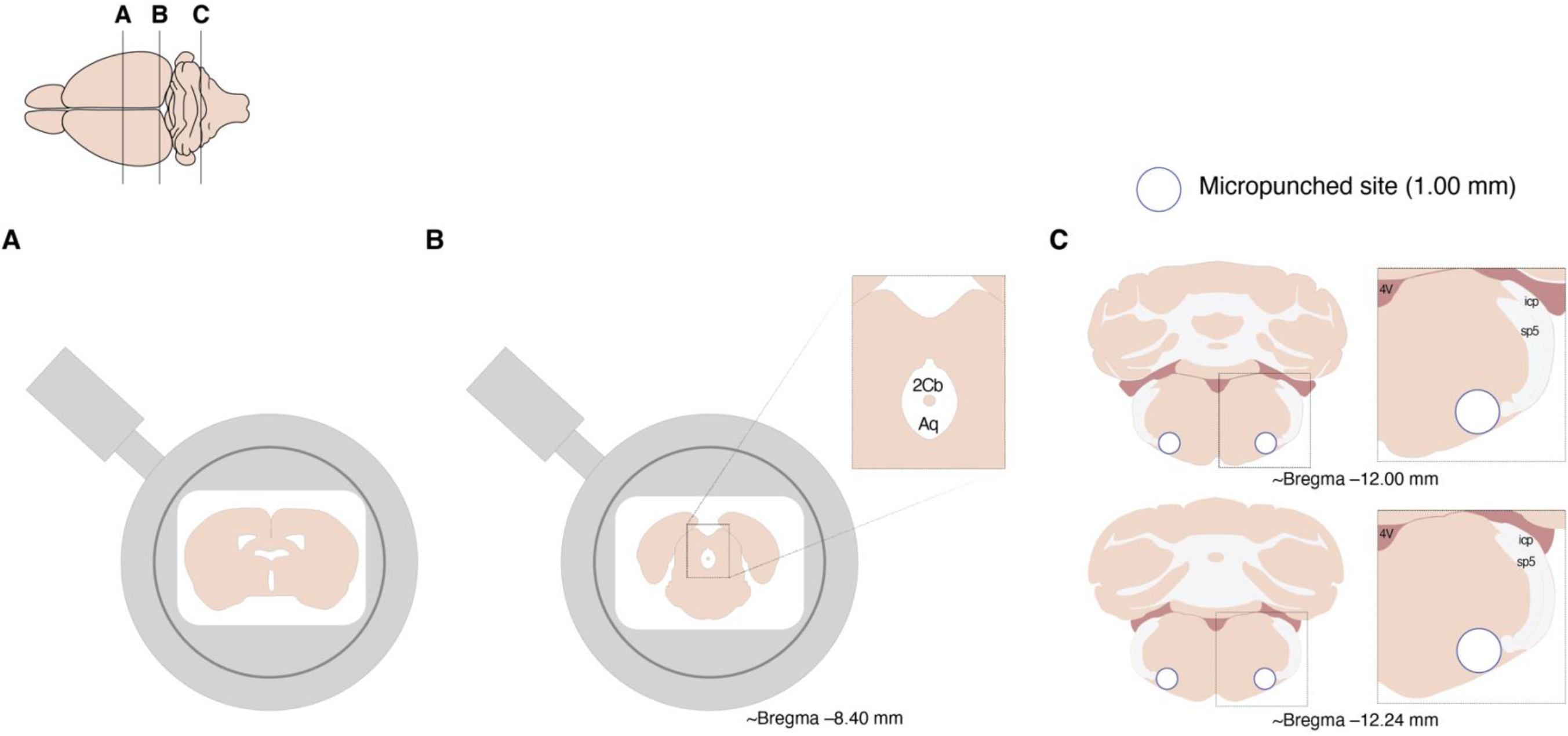
Schematic diagram of the RVLM micropunching procedure. A. Illustration of a coronally-sectioned rat brain on a cryostat sample holder after the correction of horizontal (left/right) orientation. B. Illustration of a rat brain’s block face on a cryostat sample holder displaying lobule 2 of the cerebellar vermis (2Cb) and aqueduct (Aq); inferred distance from Bregma = −8.40 mm, according to the stereotaxic coordinate system of the rat brain atlas by Paxinos & Watson (*PW7*) (Paxinos and Watson, 2014). From this point, ninety 40 μm-thick sections were trimmed rostrocaudally to access RVLM-containing sections. C. Illustration of two 200-μm thick sections and their associated inferred distances from Bregma. Micropunched RVLM-containing area is marked with blue-outlined white-filled circles. 4V; 4^th^ ventricle, icp; inferior cerebellar peduncle, sp5; spinal trigeminal tract

### RNA purification, quality, and quantity analyses

RNA from the tissue punches was extracted and purified using RNeasy Lipid tissue mini kit (Qiagen, Inc., Redwood City, CA, USA) with QIAzol Lysis reagent (Qiagen, Inc.) according to the manufacturer’s protocol. Disposable RNAase-free pellet pestles (Bio Plas, Inc., San Rafael, CA, USA) were used for the homogenization of the tissue punches. To decrease DNA contamination, columns were treated with DNase-I enzyme (Qiagen, Inc.) during the purification procedure. The quality of the RNA was analyzed using a Nanodrop Spectrophotometer (Thermo Fisher Scientific), Bioanalyzer (Agilent, Inc., Santa Clara, CA, USA) or TapeStation (Agilent, Inc.). The RNA samples used in this study have a 260/280 ratio of 1.7 ± 0.1. Seventy-seven percent of the RNA samples were analyzed for RNA integrity using either TapeStation or Bioanalyzer. High-sensitivity RNA ScreenTape System (Agilent, Inc.) was used on the TapeStation and RNA 6000 nano kit (Agilent, Inc.) was used on the Bioanalyzer. Sixty-two percent of the samples analyzed for RNA integrity had a concentration high enough to calculate RIN values; we observed a mean RNA integrity number of 7.9 ± 0.4 for these samples. The remaining 38% of the analyzed samples did not have enough concentration to calculate the RIN number; however, these RNA samples had a 28s/18s rRNA ratio of 1.8 ± 0.2. RNA concentration was estimated using either Nanodrop (73% of samples) or Qubit RNA HS assay kit (27% of samples) on a Qubit Fluorometer (Thermo Fisher Scientific).

### qPCR analysis of rat GABA and glutamate neurotransmission-related genes using RT^2^ Profiler PCR arrays

To identify the age-associated gene expression changes in GABA and glutamate neurotransmission-related genes, dsDNA binding dye (SYBR^®^ green) chemistry-based qPCRs were performed on purified RNA samples using Rat GABA & Glutamate RT^2^ Profiler PCR arrays (Qiagen, Inc., GeneGlobe ID PARN-152Z, Gene Expression Omnibus accession number-GSE220296, platform ID number-GPL32921). Each of these arrays had 96-wells. Eighty-four of the wells had primers related to GABA and glutamate neurotransmission-related genes; 44 genes were related to glutamate neurotransmission, 27 genes were related to GABA neurotransmission, and 13 genes were related to both GABA and glutamate neurotransmission. In addition, each array had five wells containing primers for the following housekeeping genes: *Actb*, *B2m*, *Hprt1*, *Ldha*, and *Rplp1*. For quality control of PCR arrays, each array had one well to assay rat genomic DNA (Rat Genomic DNA Contamination-RGDC), three wells to estimate reverse transcription efficiency (Reverse Transcription Control-RTC assays), and three wells to estimate inter-plate PCR reproducibility (Positive PCR Control-PPC assays). A list of the genes with their functional groups and well designations are reported in **Supplemental File 1**.

To perform RT^2^ profiler arrays, 50 ng of RNA collected from young (n=8), middle-aged (n=8), and aged (n=10) RVLM-containing tissue punches was reverse transcribed to cDNA using an RT^2^ preamp cDNA synthesis kit (Qiagen, Inc., First strand cDNA synthesis) following manufacturer’s guidelines, after treating the RNA samples with genomic DNA elimination buffer at 42°C for 15 min. The reverse transcription was performed using a thermal cycler (T100, Bio-Rad Laboratories, Hercules, CA, USA) at 42°C for 30 min, and at 95°C for 5 min. The first-strand cDNA synthesis step introduces external RNA control templates into the samples, and they were assayed on RT^2^ profiler arrays to estimate reverse transcription efficiency. Five μl of reverse transcribed cDNA samples were pre-amplified using RT^2^ preamp PCR master mix with PBR-152Z RT2 preamp pathway primer mix (Qiagen, Inc.) on a thermal cycler under the following conditions: 95°C (10 min) followed by 12 cycles of 95°C (15 sec) and 60°C (2 min). To eliminate the residual primers, pre-amplified samples were treated with 2 μl of side reaction reducer in a thermal cycler with the following conditions: 37°C (15 min) and 95°C (5 min). To complete the PCR array procedure, each pre-amplified sample was mixed with SYBR^®^ Green ROX qPCR master mix (Qiagen, Inc.), and 25 μl of this preparation was added into each well of the PCR array plate. Plates were covered with adhesive film, centrifuged at 3,000 rpm for 5 min, and qPCR was performed on the StepOnePlus^TM^ master cycler (Applied Biosystems^TM^, Thermo Fisher Scientific) under the following conditions; 95°C (10 min), followed by 40 cycles of 95°C (15 sec) and 60°C (1 min). At the end, melting curve analysis was performed under the following conditions 95°C (15 sec), 60°C (1 min), followed by data collection at 0.3°C increments, and 95°C (15 sec). A standard ramp rate was used for data acquisition and ROX was used as the passive reference dye.

### RT^2^ Profiler PCR arrays data analysis

To calculate threshold cycle (Ct) values, the background fluorescence of each well was identified by setting the start of baseline at cycle 3 and the end of baseline at cycle 15 for each array plate. The threshold value for Ct was manually selected and the same threshold value (1.52) was used for all the arrays. After threshold selection, Ct values of each array plate were exported as an Excel file. Each array’s quality measures, such as rat genomic DNA contamination, sample reverse transcription efficiency, and inter-plate PCR reproducibility were analyzed. If the Ct value of the RGDC well was >30 or undetermined, then the RNA sample used for that array was considered to be absent of any genomic DNA contamination. If the difference between the average of the RTC assays Ct value and the PPC assays Ct value was ≤5, then that sample was considered to have enough transcription efficiency to compare between samples. If the average of the PPC assays Ct value was 20±2, then that array was considered to be reproducible. Quality control measures for each array are reported in **Table 1**. All of the analyzed PCR arrays met the quality control measures except for one young rat (**Table 1**, Young 1503056) and one middle-aged rat (**Table 1**, Middle-aged 1504068) arrays that showed higher variance in their PCR array reproducibility measurements.

**Table 1:**
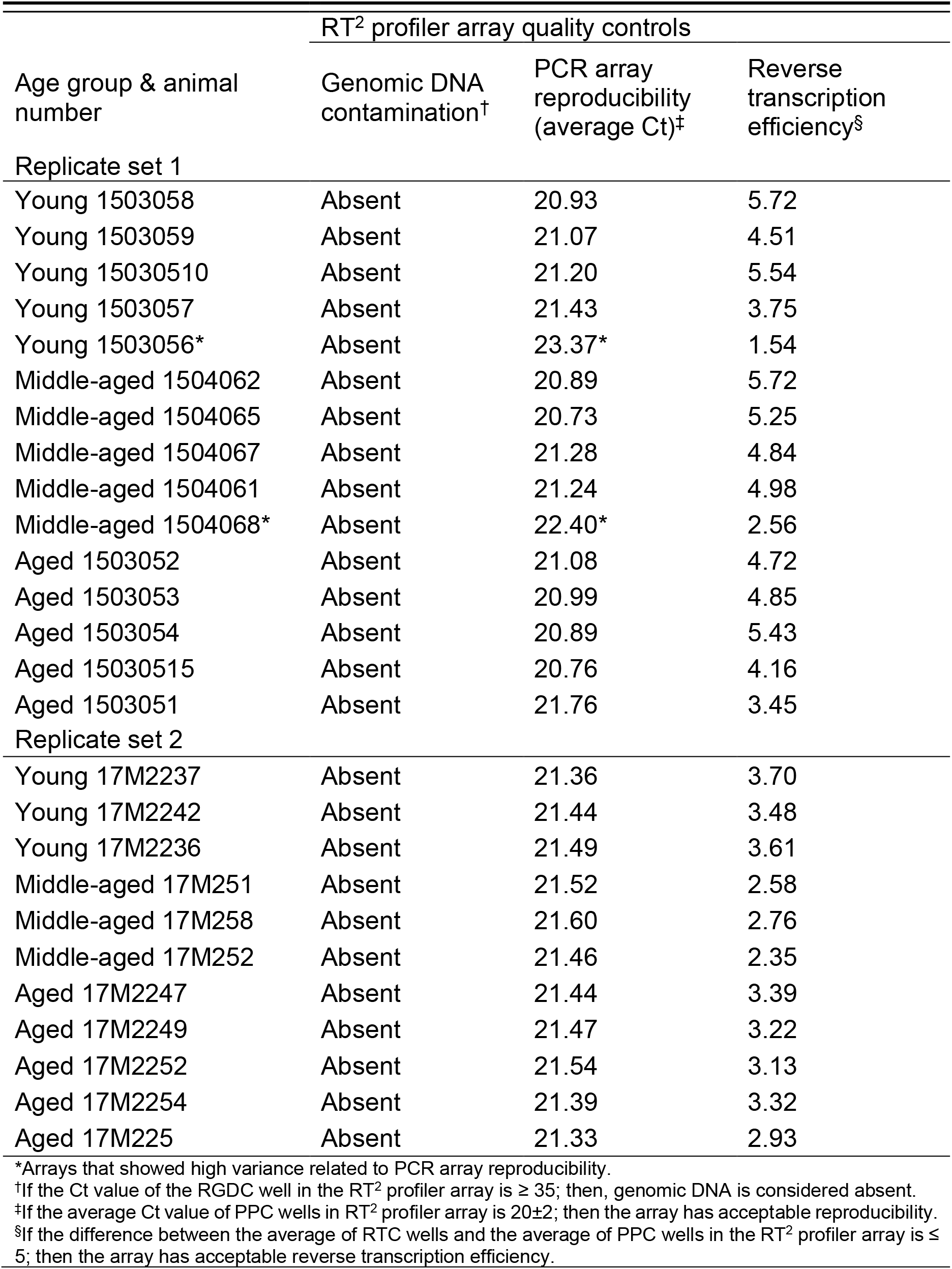
Quality control measurements of the young, middle-aged, and aged RVLM GABA and glutamate neurotransmission-related genes derived from RT^2^ profiler arrays. RGDC, Rat Genomic DNA Contamination; PPC, Positive PCR Control; RTC. Reverse Transcription Control.

| Age group & animal number | RT<sup>2</sup> profiler array quality controls |  |  |
| --- | --- | --- | --- |
|  | Genomic DNA contamination<sup>†</sup> | PCR array reproducibility (average Ct)<sup>‡</sup> | Reverse transcription efficiency<sup>§</sup> |
| Replicate set 1 |  |  |  |
| Young 1503058 | Absent | 20.93 | 5.72 |
| Young 1503059 | Absent | 21.07 | 4.51 |
| Young 15030510 | Absent | 21.20 | 5.54 |
| Young 1503057 | Absent | 21.43 | 3.75 |
| Young 1503056* | Absent | 23.37* | 1.54 |
| Middle-aged 1504062 | Absent | 20.89 | 5.72 |
| Middle-aged 1504065 | Absent | 20.73 | 5.25 |
| Middle-aged 1504067 | Absent | 21.28 | 4.84 |
| Middle-aged 1504061 | Absent | 21.24 | 4.98 |
| Middle-aged 1504068* | Absent | 22.40* | 2.56 |
| Aged 1503052 | Absent | 21.08 | 4.72 |
| Aged 1503053 | Absent | 20.99 | 4.85 |
| Aged 1503054 | Absent | 20.89 | 5.43 |
| Aged 15030515 | Absent | 20.76 | 4.16 |
| Aged 1503051 | Absent | 21.76 | 3.45 |
| Replicate set 2 |  |  |  |
| Young 17M2237 | Absent | 21.36 | 3.70 |
| Young 17M2242 | Absent | 21.44 | 3.48 |
| Young 17M2236 | Absent | 21.49 | 3.61 |
| Middle-aged 17M251 | Absent | 21.52 | 2.58 |
| Middle-aged 17M258 | Absent | 21.60 | 2.76 |
| Middle-aged 17M252 | Absent | 21.46 | 2.35 |
| Aged 17M2247 | Absent | 21.44 | 3.39 |
| Aged 17M2249 | Absent | 21.47 | 3.22 |
| Aged 17M2252 | Absent | 21.54 | 3.13 |
| Aged 17M2254 | Absent | 21.39 | 3.32 |
| Aged 17M225 | Absent | 21.33 | 2.93 |

\*Arrays that showed high variance related to PCR array reproducibility.
<sup>†</sup>If the Ct value of the RGDC well in the RT<sup>2</sup> profiler array is $\geq 35$ ; then, genomic DNA is considered absent.
<sup>‡</sup>If the average Ct value of PPC wells in RT<sup>2</sup> profiler array is $20 \pm 2$ ; then the array has acceptable reproducibility.
<sup>§</sup>If the difference between the average of RTC wells and the average of PPC wells in the RT<sup>2</sup> profiler array is $\leq 5$ ; then the array has acceptable reverse transcription efficiency.

Genes that showed a <30 Ct value in more than 50% of the arrays were used in the final analysis. Seventy-nine of the 84 GABA and glutamate neurotransmission-related genes met these criteria. In contrast, *Avp*, *Gabra6*, *Gabrr1*, *Grm6* and *Slc1a7* genes did not meet these criteria and were removed from the final analysis. Housekeeping genes that displayed less than or equal to one Ct cycle variance (Figure 2; gray-shaded regions) across arrays were considered to be consistently expressing between age groups. The Ct values for three of the five housekeeping genes (*B2m*, *Hprt1*, and *Rplp1*) were identified to be consistent between arrays (**Figure 2**) and were therefore used as reference genes for normalizing RNA expression. The geometric mean of the *B2m*, *Hprt1*, and *Rplp1* Ct values for each experimental animal/array was used for the ΔCt value calculation of each gene. A total of 79 GABA and glutamate neurotransmission-related genes’ ΔCt values were calculated. Each array/gene base 2 logarithm-transformed 2^−ΔCt^ value was used as a parameter for statistical analysis (Abdelnour, 2016). Expression changes for individual genes were considered significant if they met three criteria: fold change value >1.5 or <−1.5, t-test (unequal variance assumption) *p* value <0.05, followed by Benjamini-Hochberg False discovery rate (FDR) <0.05. Aged vs. young, aged vs. middle-aged, and middle-aged vs. young rats RVLM gene expression differences are represented as fold change values calculated using the 2^−ΔΔCt^ method (Livak and Schmittgen, 2001). All the data files containing non-normalized Ct values, normalized 2^−ΔCt^ values, and fold change values were deposited in Gene Expression Omnibus (GEO) database; GEO accession number: GSE220296, young RVLM samples: GSM6797916-GSM6797923, middle-aged RVLM samples: GSM6797924-GSM6797931, and aged RVLM samples: GSM6797932-GSM6797941.

**Figure 2:**
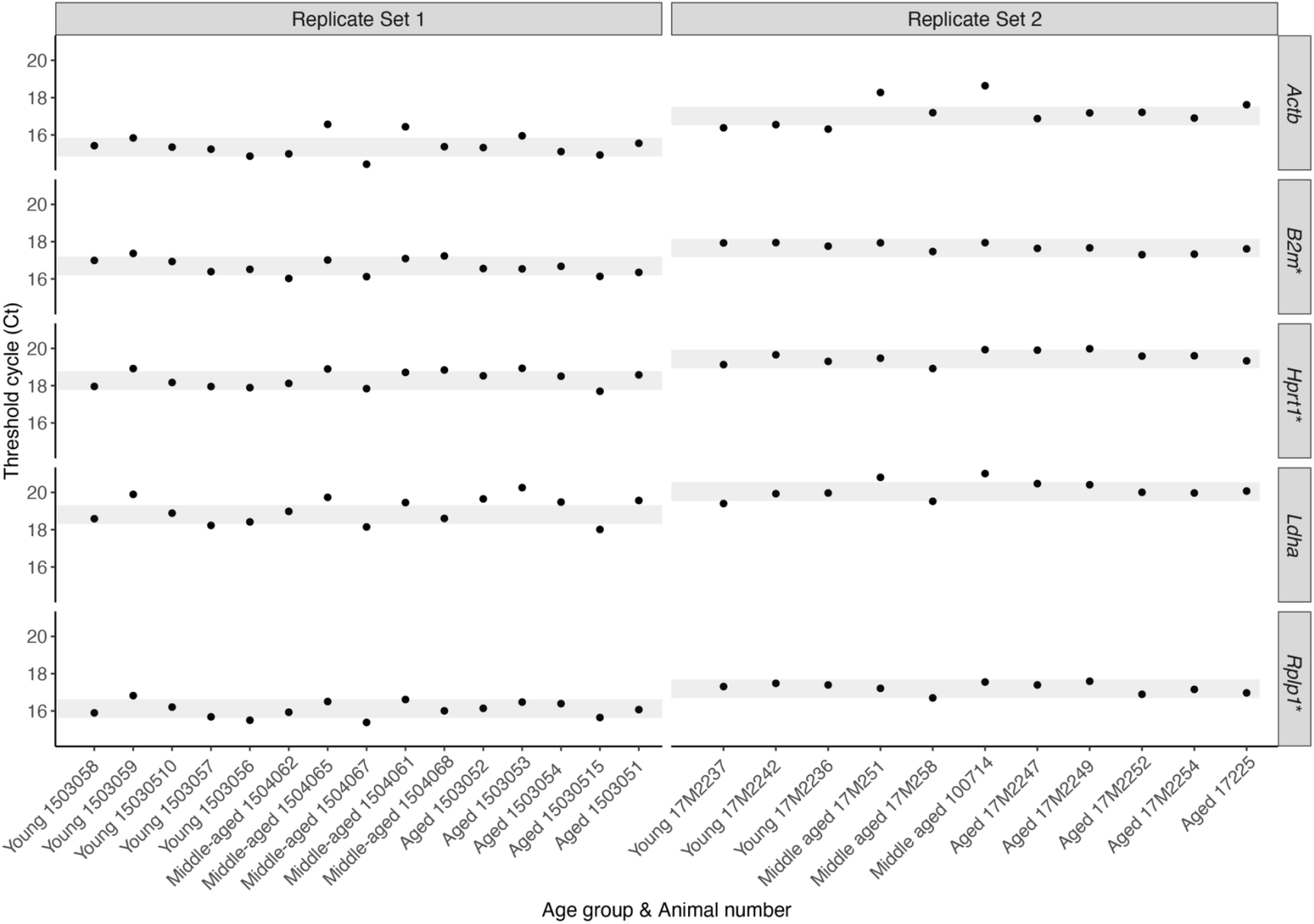
Selection of the reference genes for normalizing RNA expression in the RT^2^ profiler arrays of the young, middle-aged, and aged RVLM samples. The Ct values from the five housekeeping genes are displayed as a scatter plot for the analyzed RVLM samples. Each RVLM sample’s age group with the animal number is displayed on the x-axis, and the Ct value is displayed on the y-axis. Housekeeping genes were rendered as horizontal panels (panel names are at the right side of the figure), and replication sets were rendered as vertical panels (replication set numbers are at the top of the figure). Housekeeping genes that displayed less than or equal to one Ct cycle variance across arrays are considered to be consistently expressing between age groups. The height of gray-shaded regions indicates one Ct cycle. *B2m*, *Hprt1*, and *Rplp1* (marked with an asterisk) genes’ Ct values were inside or at the edge of the gray-shaded regions and were selected as reference genes for normalization.

### TaqMan^®^-based qPCR validation of the selected genes differential expression

The altered expression of selected genes (*Slc1a6* and *Slc6a13*) was independently validated using TaqMan^®^-based qPCR. Fifty nanograms of RNA collected from young (n=5), middle-aged (n=5), and aged (n=5) RVLM-containing tissue punches were reverse transcribed to cDNA using a High-capacity RNA to cDNA^TM^ kit (Applied Biosystems^TM^, Thermo Fisher Scientific). The reverse transcription reaction was performed using a thermal cycler (T100, Bio-Rad Laboratories) under the following conditions: 37°C (60 min), 95°C (5 min). To increase the number of qPCR reactions from each sample, cDNA was pre-amplified using TaqMan PreAmp Master Mix (Applied Biosystems^TM^, Thermo Fisher Scientific) following manufacturer’s instructions. Briefly, 5 μl of cDNA was mixed with 0.2X pooled assay mix containing *Slc1a6*, *Slc6a13*, *B2m*, *Hprt1*, and *Rplp1* genes’ TaqMan^®^ assays (Applied Biosystems^TM^, Thermo Fisher Scientific) and TaqMan^®^ PreAmp Master Mix, and preamplification was performed using a thermal cycler under the following conditions: denaturation step at 95°C for 10 min followed by 10 cycles of 95°C (15 sec) and 60°C (4 min) steps. qPCR for each gene was performed on these pre-amplified samples using TaqMan^®^ Gene Expression Master Mix on the Step One Plus master cycler (Applied Biosystems^TM^, Thermo Fisher Scientific) under the following conditions: 50°C (2 min), 95°C (10 min), followed by 40 cycles of 95°C (15 sec) and 60°C (1 min). Two technical replicates were conducted for each pre-amplified sample/gene, and reaction mixes with no template were used as negative controls. *B2m*, *Hprt1* and *Rplp1* genes were used as reference genes and the geometric mean of their Ct values was used to calculate each gene’s ΔCt value. Aged vs. young, aged vs. middle-aged, and middle-aged vs. young rats RVLM gene expression differences are represented as fold change values calculated using the 2^−ΔΔCt^ method (Livak and Schmittgen, 2001).

### Protein extraction and quantitation

To analyze proteins with immunoblotting, total proteins from RVLM-containing tissue punches were extracted using N-PER™ Neuronal Protein Extraction Reagent (Thermo Fisher Scientific) following manufacturer’s instructions. Briefly, RVLM-containing tissue punches were homogenized with disposable pestles and pellet mixer in 50 μl of 1% protease inhibitor cocktail (Sigma-Aldrich, St. Louis, MO, USA) containing N-PER™ Neuronal Protein Extraction Reagent. These homogenates were centrifuged at 10,000 × g for 10 min after incubating them on ice for 10 min. Supernatants containing protein extracts were collected. Protein concentration was estimated using biuret reaction-based Pierce BCA Protein Assay kit (Thermo Fisher Scientific). The microplate procedure recommended in the manufacturer’s user guide was followed. Incremental dilutions of bovine serum albumin (Sigma-Aldrich) were used to construct a standard curve. Colorimetric measurements were acquired on a microplate reader (Molecular Devices, Spectramax) at 562 nm. Protease inhibitor containing N-PER™ Neuronal Protein Extraction Reagent was used as a blank. Following protein quantitation, samples were aliquoted and stored at −80°C until use.

### Immunoblotting

To immunoblot protein samples, extracted proteins were first immobilized and resolved based on molecular weight on 4–15% Mini-PROTEAN TGX Precast gels (Bio-Rad Laboratories) through sodium dodecyl sulfate (SDS)-polyacrylamide gel electrophoresis (PAGE). Fifteen micrograms of protein were loaded into each well. Protein samples were denatured in 2-mercaptoethanol containing Laemmli buffer for 10 min at 95–100°C before electrophoresis. Electrophoresis was performed in a MiniProtean 4 tetra system with Tris/Glycine/SDS buffer at 90 V until the dye front reached to the bottom of the gel. Following electrophoresis, proteins immobilized on the gels were electrotransferred onto polyvinylidene difluoride membranes (Bio-Rad Laboratories, Immune-Blot PVDF Membrane) using the wet transfer method. Transfer was performed in a Mini Trans-Blot system with 20% methanol containing Tris/Glycine buffer at 200 V for 2 hr. A blue cooling unit (Bio-Rad) was used to reduce the increase in temperature.

Immunodetection for targeted proteins was conducted on PVDF membranes using the following protocol. After wet transfer, membranes were washed with deionized water three times followed by two Tris-buffered saline (TBS) washes. To decrease non-specific antibody binding, membranes were blocked for 1 hr in 5% non-fat dry milk containing TBS. After washing the membranes with 0.1% Tween-20 containing TBS for two times, membranes were incubated in primary antibody containing 0.05% Tween-20 TBS solution at 4°C overnight. Membranes were washed with 0.1% Tween-20 containing TBS for two times followed by incubation with secondary antibody for an hour. After final washes with 0.1% Tween-20 containing TBS (2 times) and TBS (4 times), membranes were imaged using SuperSignal^TM^ West Femto Maximum Sensitivity Substrate (Thermo Fisher Scientific) following manufacturer’s guidelines. A Kodak Image Station 4000 was used to capture the images of the chemiluminescence bands. Each wash was performed for 10 min. Information on the primary and secondary antibodies used in this study are reported in **Table 2**. The anti-EAAT4 antibody we used was validated using proteins purified from positive and negative tissue controls. Proteins purified from liver and cerebellum samples were used as negative and positive controls, respectively.

**Table 2:** List of the antibodies used for immunoblot analyses of EAAT4 and β-actin proteins.

| Antibody | Host | Clonality | Source | Catalog number | Lot number | RRID | Titer used for immunoblotting |
| --- | --- | --- | --- | --- | --- | --- | --- |
| Anti-EAAT4, primary | Rabbit | Recombinant monoclonal IgG | Abcam | ab186435 | GR161408-1 | AB_2868394 | 1:500 |
| Anti-β-actin, primary | Mouse | Monoclonal IgG1 [AC-15] | Abcam | ab6276 | GR231981-1 | AB_2223210 | 1:5000 |
| Anti-rabbit HRP tagged, secondary | Goat | Polyclonal IgG, H&L | Abcam | ab6721 | GR208028-2 | AB_955447 | 1:5000 |
| Anti-mouse peroxidase tagged, secondary | Goat | Polyclonal IgG | Sigma-Aldrich | A4416 | SLBK1590V | AB_258167 | 1:2000 |

### Nissl staining & Microscopy

Archived (at –80°C) young (n=9, 18 sections containing a total of 36 micropunched sites), middle-aged (n=11, 18 sections containing a total of 36 micropunched sites), and aged (n=8, 16 sections containing a total of 32 micropunched sites) 200 μm-thick coronal sections that had their RVLM-containing area micropunched were stained. These animals were independent from the ones used for RNA and protein analyses. Sections were thawed on charged glass slides at room temperature and dried overnight in an oven at 40°C. After drying, sections were cleared with 1:1 chloroform (Fisher Scientific): ethyl alcohol (Fisher Scientific) solution for four days to remove lipids from the sections. Following lipid clearing, slides were treated with xylenes (Sigma-Aldrich) twice for 10 min each. For the purpose of aqueous-based staining, slide-mounted sections were rehydrated through descending ethanol concentrations (100%, 95%, 70%, 50%, and deionized water; 3 min each). These sections were stained overnight in a solution containing 0.012% Cresyl Violet (Sigma-Aldrich) in acetate buffer (pH 3.5) at 40°C in an oven. Following overnight staining, sections were washed with running water (3 min) and dehydrated through ascending ethanol concentrations (deionized water, 50%, 70%, 95%, and 100%; 3 min each). Dehydrated sections were treated with xylenes (twice for 15 min each), coverslipped with DPX mountant (Sigma-Aldrich), and air-dried inside a fume hood. Large field-of-view images of the stained sections were captured with a motorized Olympus BX63 upright microscope mounted with a DP74 color camera using a Plan Apochromatic ×10 magnification objective lens (0.40 numerical aperture) (Olympus). Images were acquired and stitched using cellSens software (Olympus), and exported as TIFF-formatted files.

### Standardized atlas-based mapping of the micropunched sites

Micropunched sites were mapped onto the *Brain Maps 4.0 (BM4.0)* rat brain reference atlas (Swanson, 2018) following the method described previously (Khan et al., 2018). The photomicrographs of stained sections were evaluated by laboratory members using the cytoarchitectural descriptions outlined in *BM4.0* to determine the corresponding atlas level of the micropunched ventrolateral medullary area. Due to dorsoventral discrepancies in the plane of section of the tissue analyzed, the area in close proximity to the micropunched site was used in deducing the atlas level. The images were imported into Adobe Illustrator (AI; Version 26, Adobe Inc., San Jose, CA, USA), where the cytoarchitecture revealed by Nissl staining permitted the delineation of gray matter regions and white matter tracts using the *Pencil* tool in a separate layer in AI. Following brain region parcellation, the digital template of the corresponding *BM4.0* atlas level was imported and the layers containing the section image and traced regional boundaries were transposed to accurately align with the atlas level template for precise localization of the sampled site. To map all micropunched sites to the right-side of the atlas template for ease of comparison, the image was mirrored vertically when mapping the left-side micropunched site. Once aligned, the micropunched site was traced in a new layer in AI. If the area surrounding the micropunched site was damaged, distorting the bounds of the punch, the micropunched site was not mapped (Young = 7, Middle-aged = 7, Aged = 0). If the tissue near the micropunched site was subject to “hanging”, or was displaced when mounted, the outline of the punch was drawn as if the position of the hanging portion had not been altered. Tracings of the micropunched sites that coincided to the same atlas levels were overlaid without modification of their two-dimensional spatial location on the template. The spatial distribution of these tracings was then analyzed further as described in the next section.

### Density analysis of the sampled sites

Sampled sites were analyzed in one (anterior-posterior) and two (dorsoventral and mediolateral) dimensions. To analyze anterior-posterior distribution of the micropunch sites, the number of sampled site tracings mapped onto each atlas level was counted for each age group. Anterior-posterior cumulative density probabilities of the sampled sites for each age group was visualized using RStudio *ggplot2* package (Wickham, 2016) with empirical cumulative density function. Differences in probability distribution between age groups were tested using the non-parametric Kolmogorov-Smirnov test (two sample K-S test) and a *p* value <0.05 was considered as statistically significant.

To analyze dorsoventral and mediolateral density distributions of the sampled sites, each sampled tracing was filled with red color at 10% opacity. Overlaid sampled site traces from each level for each age group were exported as PNG files. Bounding box size (1135 × 799 pixels) and resolution (300 ppi) were consistent for all the exports. PNG images were imported into RStudio and pixel values were color coded using functions from *imager* (Barthelme, 2021) and *spatstat* (Baddeley et al., 2015) packages. Briefly, each PNG image was processed using Gaussian smoothing (sigma = 5) and converted into a grayscale image. These grayscale images were transformed into *spatstat* objects and pixel values were color coded. Low and high pixel values were interpreted as high and low sampled site densities, respectively. Color coding parameters were normalized for each age group to spatially visualize relative sampled site density. Each color-coded image was exported as an SVG file with the same size bounding box as the PNG image and overlaid on the top of the corresponding atlas level template.

### Statistical and descriptive analyses

Normalized and scaled gene expression values of the arrays were analyzed using Principal Component Analysis (PCA). *Stats* and *rgl* (Adler et al., 2020) packages in RStudio (RCoreTeam, 2019) were used for the analysis and plotting. Differentially expressed genes identified in aged vs. young, middle-aged vs. young, and aged vs. middle-aged RT^2^ Profiler array comparisons were considered significant if they met the following criteria: fold change value >1.5 or < −1.5, independent t-test (unequal variance assumption) with a *p* value <0.05, followed by Benjamini-Hochberg False discovery rate correction (FDR) with a *p* value <0.05. Volcano plots and their density contours, Euler diagram, heatmaps, density curves and boxplots were graphed using *eulerr* and *ggplot2* package (Wickham, 2016) in RStudio. Density values of the immunoblot bands were measured using ImageStudio Lite software (LI-COR Biotechnology, Lincoln, NE, USA). Normalized density values were compared using ANOVA followed by Tukey’s High Significance Difference *post hoc* test, and a *p* value <0.05 was considered as statistically significant. All the statistical tests were performed in RStudio (RCoreTeam, 2019).

## Results

### RT^2^ Profiler PCR Array analysis of GABA and glutamate neurotransmission-related gene expression in the RVLM

To visualize relationships between young (n=8), middle-aged (n=8), and aged (n=10) RVLM RT^2^ Profiler PCR arrays, ΔCt values were analyzed using Principal Component Analysis (PCA). The first three principal components calculated in the PCA analysis explain 86% of the variation in the analyzed arrays (Principal Component 1: 65.2%; Principal Component 2: 15.6%; and Principal Component 3: 5.2%) and results are plotted as a three-dimensional scatter plot in **Figure 3**. Most of the middle-aged (**Fig. 3**, *orange spheres*) and aged arrays (**Fig. 3**, *red spheres*) showed a spatial separation from the young arrays (**Fig. 3**, *green spheres*), whereas middle-aged arrays were not spatially separated from aged arrays (**Fig. 3**).

**Figure 3:**
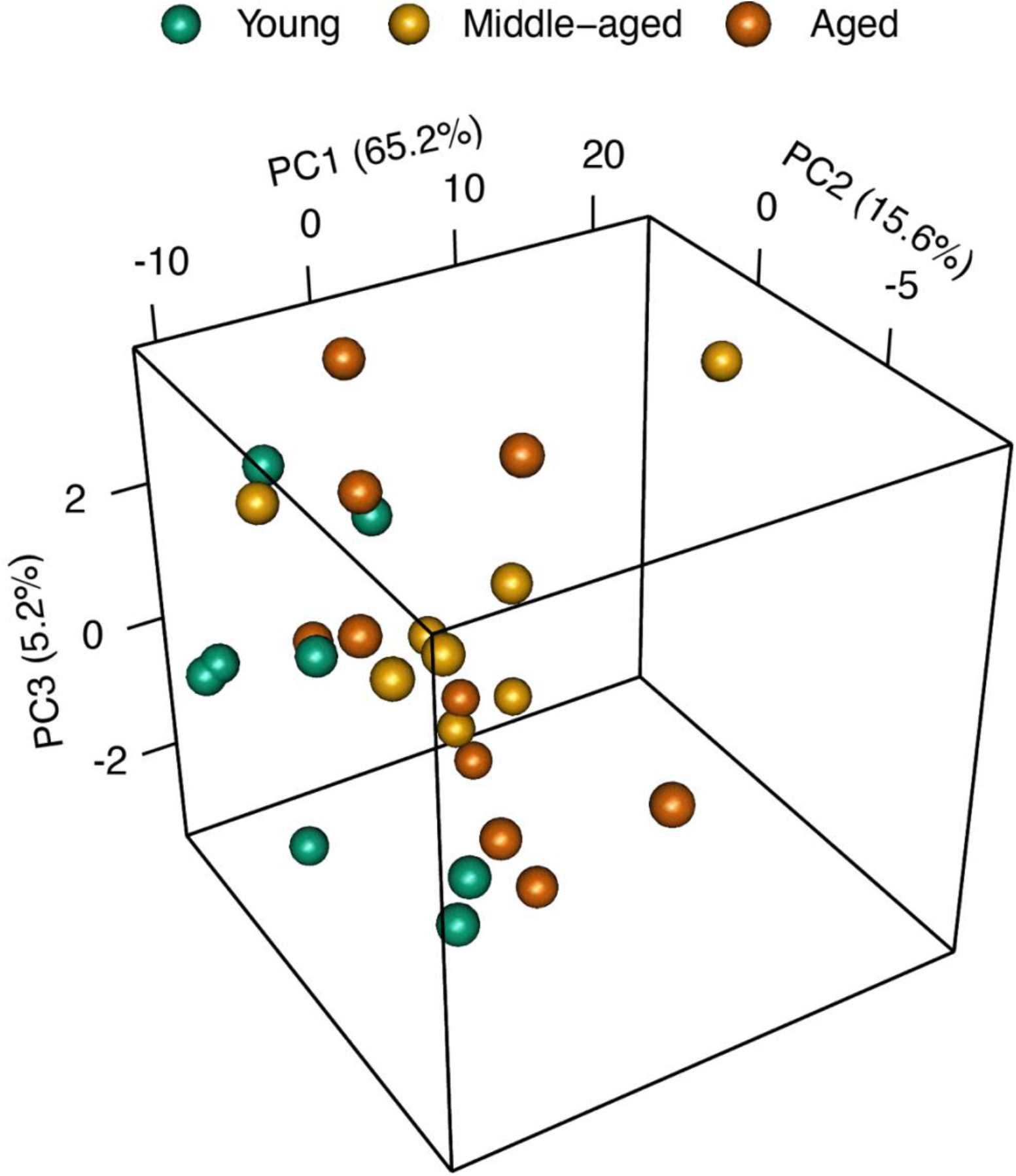
Principal Component Analysis of GABA and glutamate neurotransmission-related gene expression. The ΔCt values from the RT^2^ Profiler PCR arrays were analyzed with Principal Component Analysis, and the first three principal components were plotted as a three-dimensional scatter plot. Each array is represented as a sphere and color-coded based on the age group. Sample size for young = 8, middle-aged = 8, and aged =10.

To identify age-associated, differentially expressed GABA and glutamate neurotransmission-related genes in the RVLM, fold change values were calculated between age groups for the sorted 79 genes. Gene fold change values (log2 transformed) are plotted against their p-values (−log10 transformed) as volcano plots in **Figure 4A**, and as two-dimensional density plots in **Figure 4B**, for aged vs. young (**Fig. 4** *top*), middle-aged vs. young (**Fig. 4** *middle*), and aged vs. middle-aged (**Fig. 4** *bottom*) comparisons. Twenty-three of the 79 genes showed significant altered RVLM expression in aged compared to young rats (**Fig. 4A** *top*, *red dots*, |fold change| >1.5, t-test *p* < 0.05, FDR *p* < 0.05). All of these genes were identified to be downregulated and their absolute fold change values were less than two, with the *Slc1a6* gene demonstrating the highest fold downregulation. In addition, most of the tested genes demonstrated a downregulatory trend in the aged RVLM (**Fig. 4A** *top*) which is more evident when the aged vs. young volcano plot is rendered as a two-dimensional density plot in **Fig. 4B** *top*. As displayed in the aged vs. young comparison in **Fig. 4B**, the highest density of gene dots is on the negative side of the fold change x-axis (**Fig. 4B** *top*, *light gray* contours representing high relative density).

**Figure 4:**
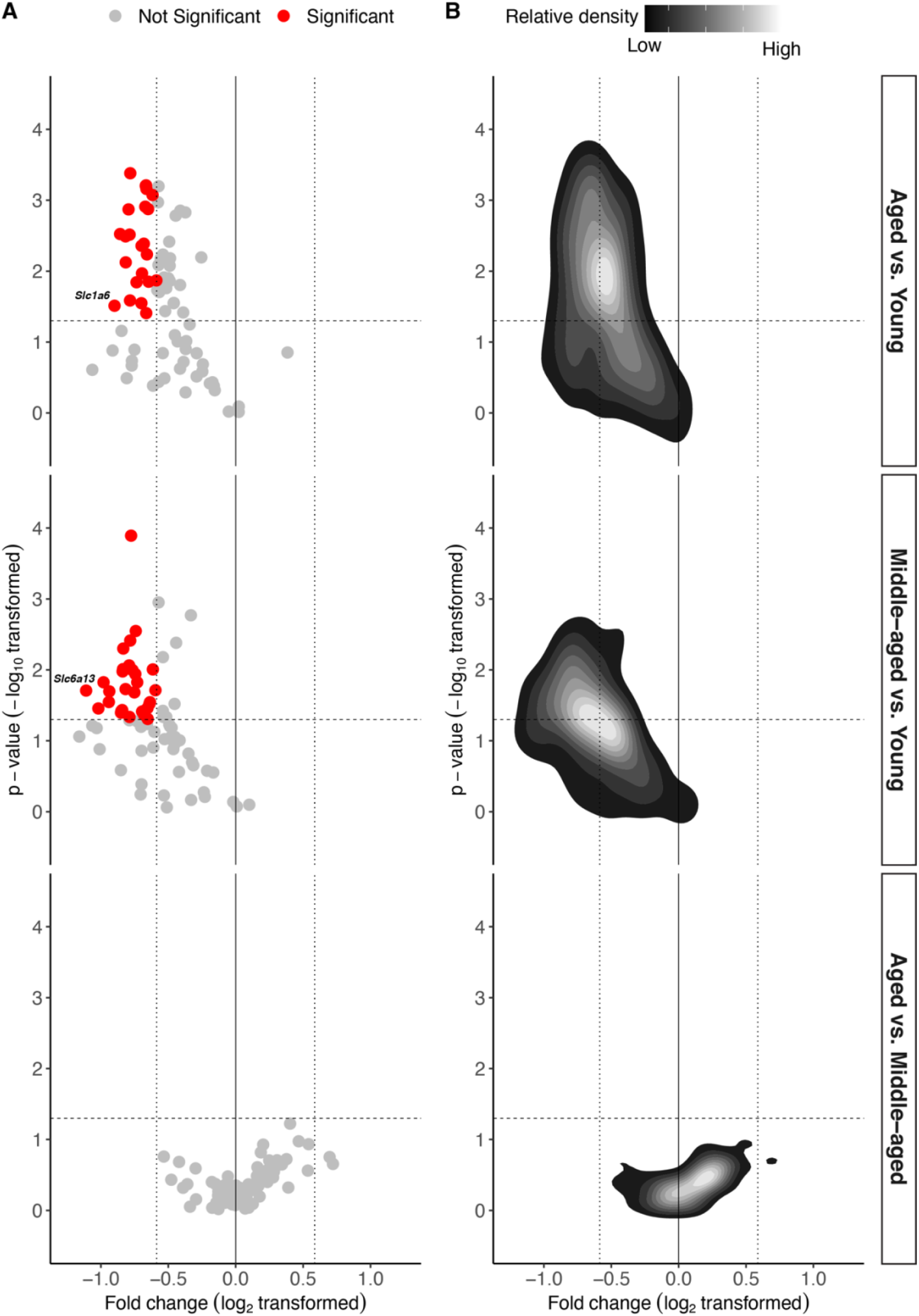
The majority of GABA and glutamate neurotransmission-related genes displayed downregulation at the mRNA level in the RVLM of middle-aged and aged rats. **A.** Age-associated GABA and glutamate neurotransmission-related gene expression changes in the RVLM are presented as volcano plots. Genes are represented as points, and statistically significant genes that showed 1.5-fold change are coded in *red*. **B.** Point patterns in A are displayed as two-dimensional density plots with high and low densities colored as *gray* and *black* contours, respectively. In A & B, plots are paneled horizontally based on age group comparisons, and panel labels are on the figure’s right side. Statistical significance: t-test *p* < 0.05 with < 0.05 FDR, and is marked with a horizontal dashed line. 1.5-fold change is marked with vertical dotted line. Sample size for young = 8, middle-aged = 8, and aged = 10.

Thirty of the 79 GABA and glutamate neurotransmission-related genes showed significantly-altered expression in the RVLM of middle-aged compared to young rats (**Fig. 4A** *middle*, *red dots*, |fold change| >1.5, t-test *p* < 0.05, FDR *p* < 0.05). All of these genes were identified to be downregulated and the *Slc6a13* gene showed the highest fold downregulation. Similar to the aged vs. young comparison, a downregulatory trend was observed when the volcano plot is rendered as a two-dimensional density plot in middle-aged vs. young comparison (**Fig. 4B** *middle*, *light gray* contours representing high relative density). There were no differences in gene expression in aged vs. middle-aged comparisons (**Fig. 4B** *bottom*).

An Euler diagram was used to show relationships between downregulated GABA and glutamate neurotransmission-related genes identified in aged vs. young and middle-aged vs. young comparisons (**Figure 5**). Twelve of the same genes were identified to be downregulated in aged vs. young and middle-aged vs. young comparisons (**Fig. 5**, *taupe* intersection).

**Figure 5:**
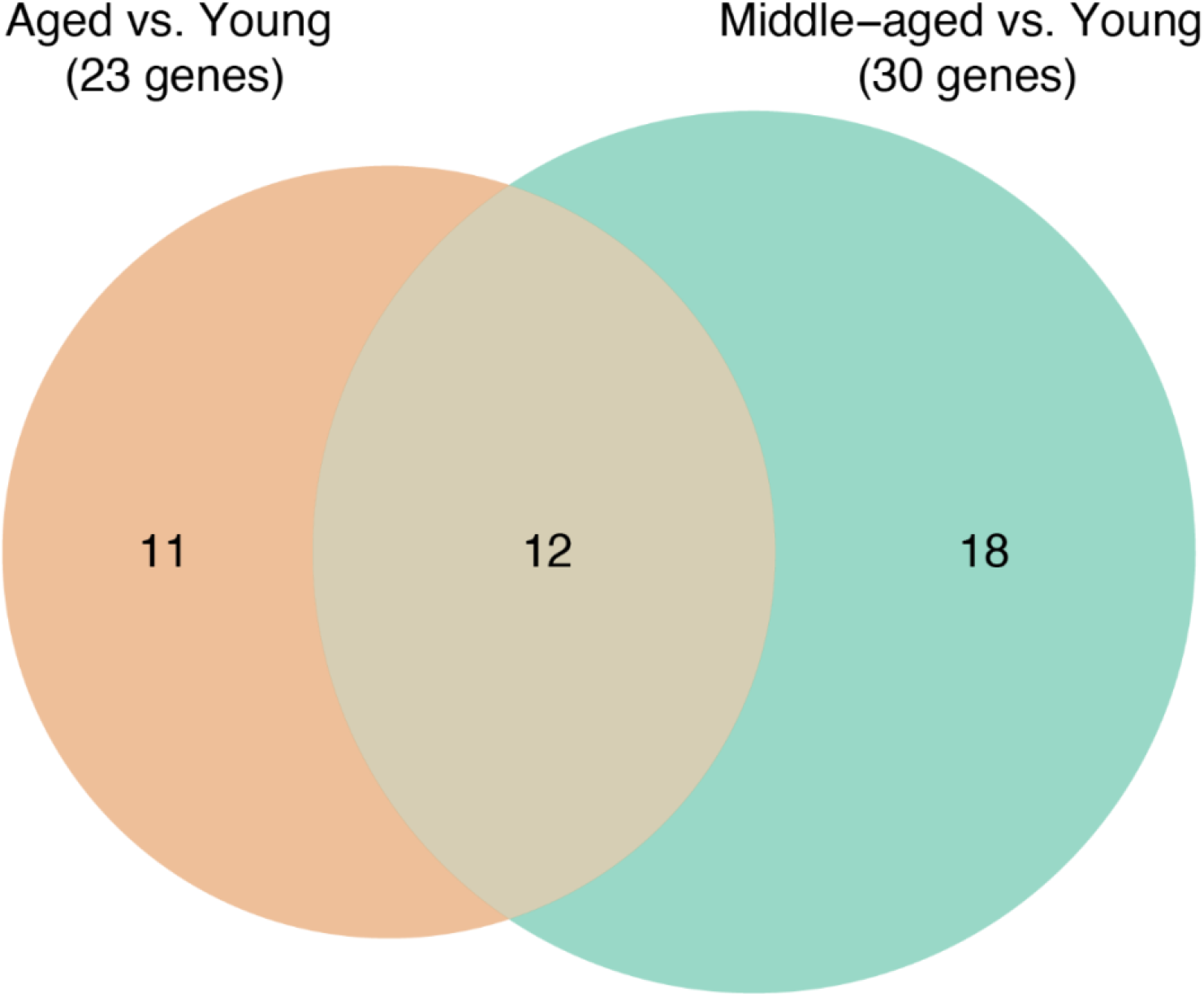
GABA and glutamate neurotransmission-related gene expression at the mRNA level was decreased by middle age in the RVLM of F344 rats. The relation between the genes that showed statistically significant differences with >1.5 fold decreases in aged vs. young and middle-aged vs. young comparisons are represented with an Euler diagram. Twelve common genes were down-regulated in both middle-aged and aged compared to the RVLM of young rats. Statistical significance: t-test *p* < 0.05 with < 0.05 FDR.

To identify the major functional category of the downregulated genes, expression of the sorted (fold change < −1.5, t-test *p* < 0.05, FDR *p* < 0.05) GABA and glutamate neurotransmission-related genes, and their annotated functional categories, were displayed as heat maps in **Figure 6**. Genes identified in the aged vs. young and middle-aged vs. young comparisons were visualized in **Figure 6A** and **B**, respectively. The annotated functional category and sub-categories of the genes are displayed at the left side of **Fig. 6A** and **B** heat maps. Of the sorted genes, 65.2% in the aged vs. young comparisons were related to glutamate neurotransmission (**Fig. 6A**, Glutamatergic cluster), and 60% in the middle-aged vs. young comparisons were related to glutamate neurotransmission (**Fig. 6B**, Glutamatergic cluster). Of the twelve common genes identified in the aged vs. young and middle-aged vs. young comparisons (**Fig. 6A** & **B**, marked with an asterisk), 9 (75%) were related to glutamate neurotransmission.

**Figure 6:**
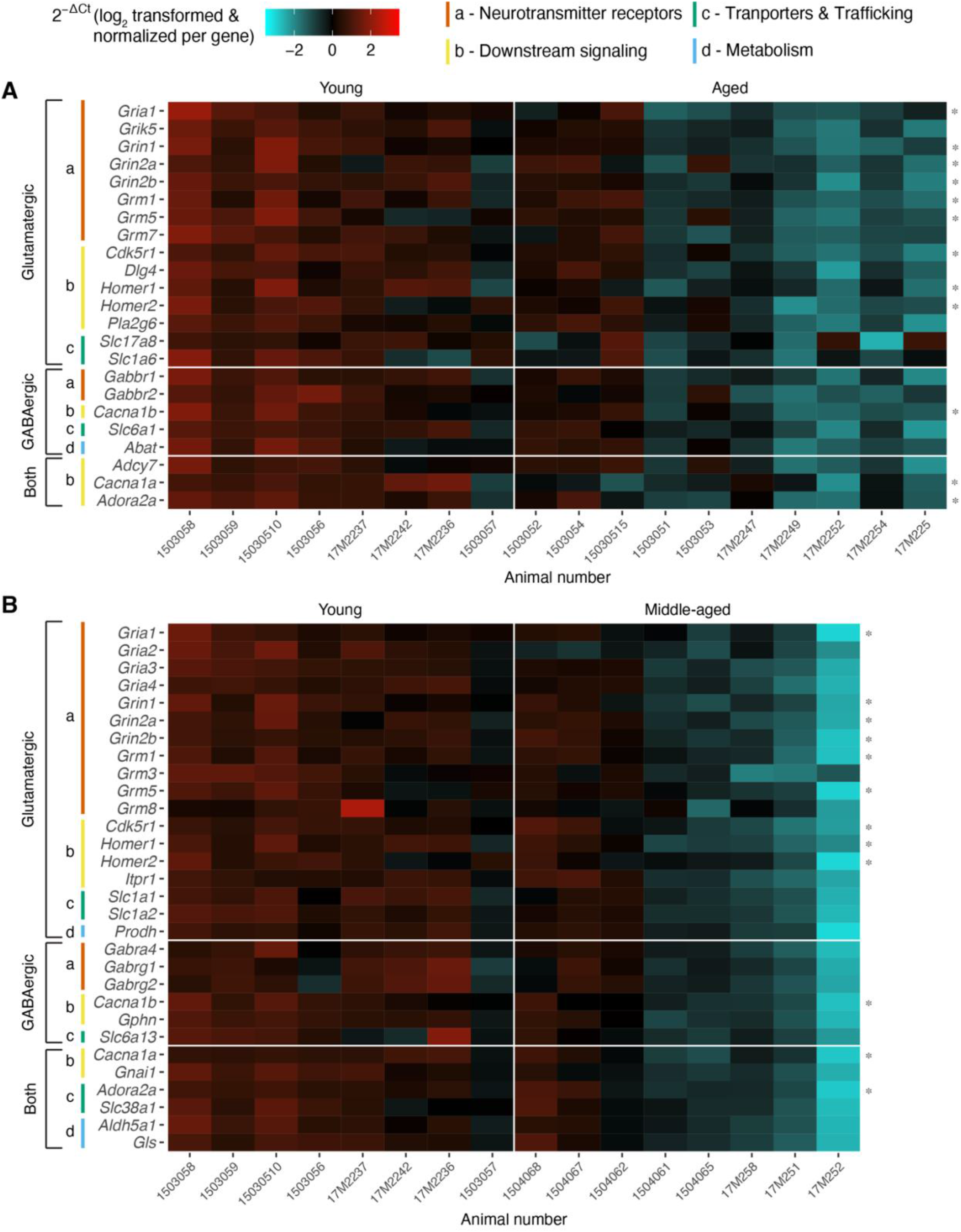
Heat map visualization of the age-associated downregulation of GABA and glutamate neurotransmission-related genes. Gene expression values (2^—ΔCt^) of the differentially downregulated genes (t-test *p* < 0.05 with *p* < 0.05 FDR, < 1.5 fold change) are represented as heat maps in A for the aged vs. young comparison, and in B for the middle-aged vs. young comparison. Animal numbers and gene symbols are represented on the x and y axes, respectively. The age category of the animals is presented at the top of each heat map. For the visualization purpose, 2^−ΔCt^ values of each gene were normalized separately, and the scale bar for the colors is at the top of the figure. Annotated functional categories and sub-categories of the genes are marked at the left side of the heat maps. Functional sub-categories are coded with color and letter notation, and the key is at the top of the figure.

### Effect of the reference genes selection on the age-associated GABA and glutamate neurotransmission-related gene expression in the RVLM

To identify the effect of reference gene selection on the observed downregulatory trend of GABA and glutamate neurotransmission-related genes in the aged vs. young and middle-aged vs. young RVLM comparisons, RT^2^ profiler PCR arrays data were analyzed at the Ct value level, and after normalizing the Ct values with different combinations (sets of two reference genes) of the selected three reference genes. Ct value differences were compared between age groups, and the density curves of the Ct value differences are displayed in **Figure 7A**. Positive and negative Ct value differences imply downregulatory and upregulatory trends, respectively. Genes that displayed positive Ct value differences (**Fig. 7A** *bottom red marks*) demonstrated more area under the density curve (**Fig. 7A**, *gray area under the curve*) than the genes that displayed negative Ct value difference (**Fig. 7A**, *white area under the curve*). Density curves of the analyzed genes fold change values (log2 transformed) calculated using *B2m* and *Hprt1*, *B2m* and *Rplp1, and Hprt1* and *Rplp1* reference gene combinations were displayed in **Figure 7B**. Genes that showed negative fold change values (**Fig. 7B** *bottom red marks*) have more area under the density curve (**Fig. 7B**, *gray area under the curve*) than the genes that displayed positive fold change value (**Fig. 7B**, *white area under the curve* and *green marks at the bottom*). For direct comparison, density curves of the fold change values calculated using the geometric mean of *B2m*, *Hprt1*, and *Rplp1* reference genes were displayed in **Fig. 7B** top panel.

**Figure 7:**
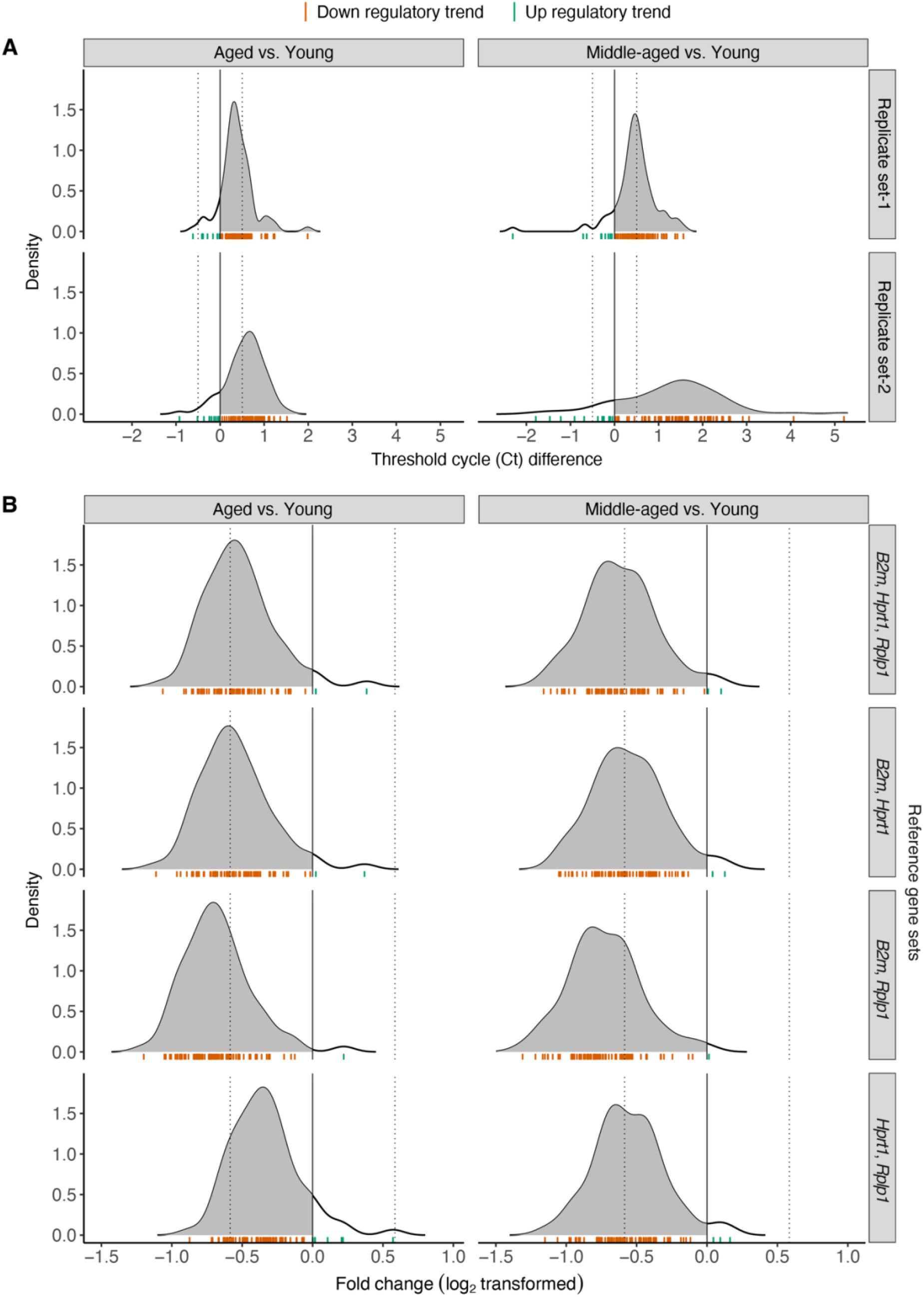
Selection of the reference genes did not affect the GABA and glutamate neurotransmission-related genes down-regulatory trend observed in the aged and middle-aged RVLM arrays. A. Density plots of the GABA and glutamate neurotransmission-related genes Ct value differences for aged vs. young and middle-aged vs. young comparisons. Plots are paneled horizontally based on replication sets, and panel labels are on the figure’s right side. Sample size for replicate set 1; young n=5, middle-aged n=5, and aged n=5. Sample size for replicate set 2; young n=3, middle-aged n=3, and aged n=5. B. Density plots of the GABA and glutamate neurotransmission-related genes fold change values (log_2_ transformed) for the aged vs. young and middle-aged vs. young comparisons. Plots are paneled horizontally based on the reference gene sets used for the fold change values calculation. Sample size for young n=8, middle-age n=8, and aged n=10. A and B are paneled vertically based on age group comparisons, and panel labels are at the top of each Figure. Gray-colored area under the curve in the density plots represents the density of the genes that showed down regulatory trend. In addition, genes were represented as marks at the bottom of each density plot. Red and green marks represent genes that showed down-regulatory and up-regulatory trends, respectively.

### TaqMan^®^ real-time PCR validation of the age-associated downregulation of the genes Slc1a6 and Slc6a13 in the RVLM

Of the analyzed genes, *Slc1a6* and *Slc6a13* displayed the highest fold downregulation in the RVLM of aged and middle-aged rats, respectively (**Fig. 4**). To validate the age-associated differential expression of *Slc1a6* and *Slc6a13*, RNA extracted from young (n=5), middle-aged (n=5), and aged (n=5) RVLM samples were analyzed using TaqMan^®^ real-time PCR. The fold change values of *Slc1a6* and *Slc6a13* genes were calculated from the aged vs. young, middle-aged vs. young, and aged vs. middle-aged comparisons using RT^2^ Profiler array and TaqMan^®^ real-time PCR and are presented in **Table 3**. Similar to the RT^2^ Profiler array analysis, in TaqMan^®^ real-time PCR, *Slc1a6* gene expression was observed to be down-regulated more than 1.5 fold in the aged compared to the young RVLM (**Table 3**). *Slc6a13* also showed a similar trend for the middle-aged vs. young comparison (**Table 3**).

**Table 3:**
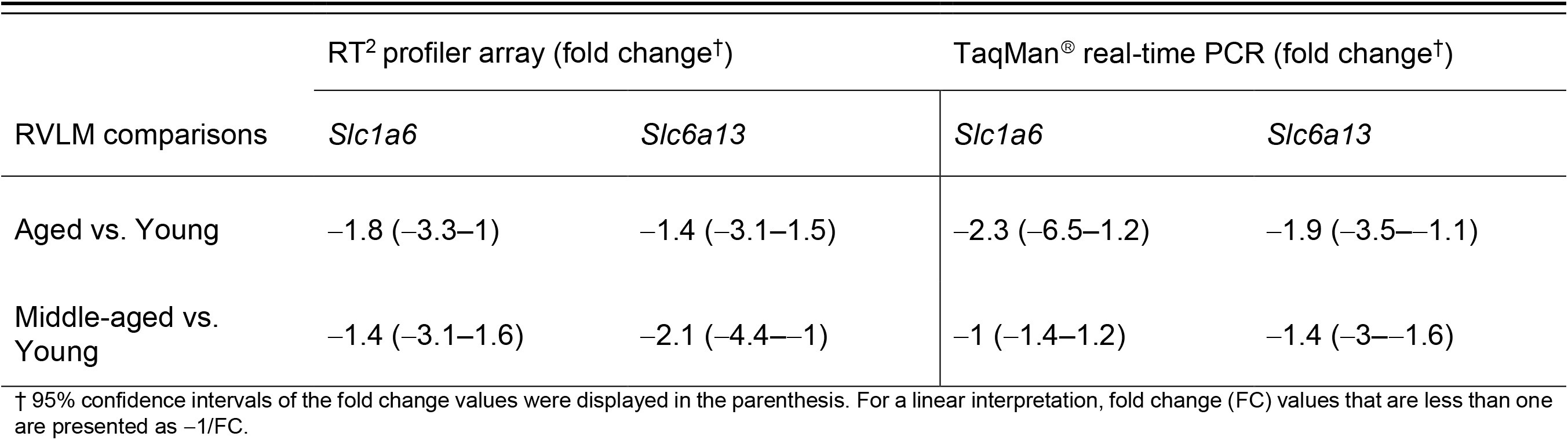
Independent validation of the age-associated *Slc1a6* and *Slc6a13* gene downregulation in the RVLM using TaqMan^®^ real-time PCR.

### Immunoblot analysis of EAAT4 (Slc1a6) protein expression

To validate the age-associated downregulation of the *Slc1a6* gene observed in the aged vs. young comparison, protein expression of EAAT4 was analyzed using the immunoblot method on the young, middle-aged, and aged RVLM protein samples. First, anti-EAAT4 antibody specificity was validated with positive and negative tissue controls. Chemiluminescence bands resulting from the anti-EAAT4 antibody reactivity on rat cerebellum, liver, and RVLM protein samples were displayed in **Figure 8A**. Protein samples purified from the cerebellum were used as positive controls, and the liver protein sample was used as a negative control. As shown in **Fig. 8A**, the anti-EAAT4 antibody reacted to a protein between 50 and 75 KDa (marked with an asterisk, predicted M.W. ∼65KDa) in the cerebellum and RVLM lanes. In the same position, no band was observed in the liver protein lane. In addition, the EAAT4 antibody displayed non-specificity by reacting to proteins, which are either isoforms of EAAT4 or other proteins (**Fig 8A**, bands other than the ones in the row marked with an asterisk).

**Figure 8:**
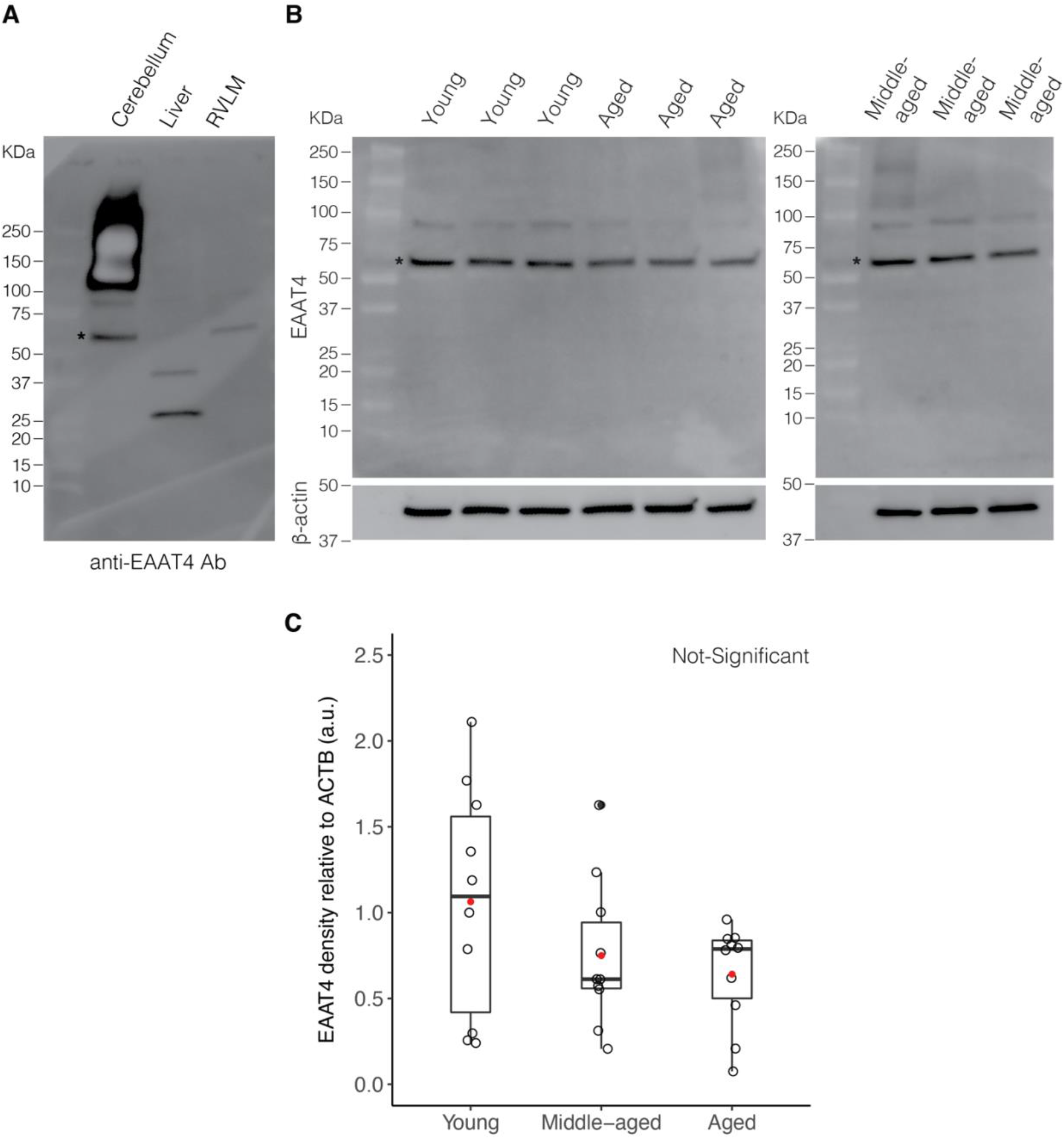
Immunoblot analysis of the young, middle-aged, and aged RVLM protein samples for Excitatory amino acid transporter 4 (EAAT4) protein expression changes. A. Immunoblot showing anti-EAAT4 antibody (Abcam, Cat#ab186435) reactivity on rat cerebellum, liver, and RVLM protein samples. Cerebellum and liver protein samples were used as positive and negative controls for EAAT4 protein expression. The row containing EAAT4 (predicted M.W. ∼65 kDa) bands is marked with an asterisk. B. Proteins purified from the young, middle-aged, and aged RVLM micropunches were blotted with an anti-EAAT4 antibody and the resulting chemiluminescence bands are displayed. EAAT4 (predicted M.W. ∼ 65 kDa) bands containing row is marked with an asterisk. β-actin (predicted M.W. ∼ 42 kDa) protein expression was used for the normalization of EAAT4 protein quantitation and their chemiluminescence bands are shown at the bottom. C. The density of EAAT4 (n=10) protein bands relative to that of β-actin bands are plotted as box plots. Filled red dots represent average density values of each age group.

The proteins purified from RVLM micropunches in young, middle-aged, and aged rats were blotted with the anti-EAAT4 antibody, and the chemiluminescence bands are presented in **Figure 8B**. Beta-actin (β-actin) expression (**Fig. 8B** *bottom*) was used for the normalization of EAAT4 protein expression. EAAT4 chemiluminescence bands displayed a lower density trend in the aged compared to young RVLM samples (**Fig. 8B**, marked with an asterisk, predicted M.W. ∼65 KDa). Semi-quantitative densitometry analysis of the normalized EAAT4 bands’ intensity confirmed this downregulatory trend in the RVLM of aged rats (**Figure 8C**). In addition, this downregulatory trend was also observed in the middle-aged RVLM (**Fig. 8C**). However, the differences were not statistically significant (**Fig. 8C**, n=10 for each age group, ANOVA *p* < 0.05 followed by Tukey HSD *p* < 0.05).

### Validation of the RVLM micropunching method through sampled site mapping

To validate the RVLM micropunching method and the anatomical location of the sampled site, micropunched sites from young (n=9, 29 micropunch sites), middle-aged (n=11, 29 micropunch sites), and aged (n=8, 32 micropunch sites) rats were mapped onto Brain Maps 4.0 (*BM4.0*) rat brain reference atlas templates (Swanson, 2018). Representative images of the stained sections containing RVLM micropunched sites are presented for each age group in **Figure 9** (dashed lines-gray matter region boundaries, solid lines-white matter tracts). Outline of the micropunch-sampled site was identified with a red color trace on each section (**Fig. 9**). Maps containing micropunch-sampled sites for young, middle-aged and aged groups at each level are presented in **Figure 10**, and each red trace represents one micropunch-sampled site. All the sampled sites were observed on the ventrolateral side of the atlas templates (**Figure 10**). Micropunched sections from young rats were determined to correspond to *BM4.0* atlas levels 56–65 (**Figure 10**, Young, left panel). In the middle-aged group, the micropunched sections corresponded to atlas levels 57, and 59–65 (**Figure 10**, Middle-aged, middle panel). Sections from aged rats corresponded to atlas levels 58–68 (**Figure 10**, Aged, right panel).

**Figure 9:**
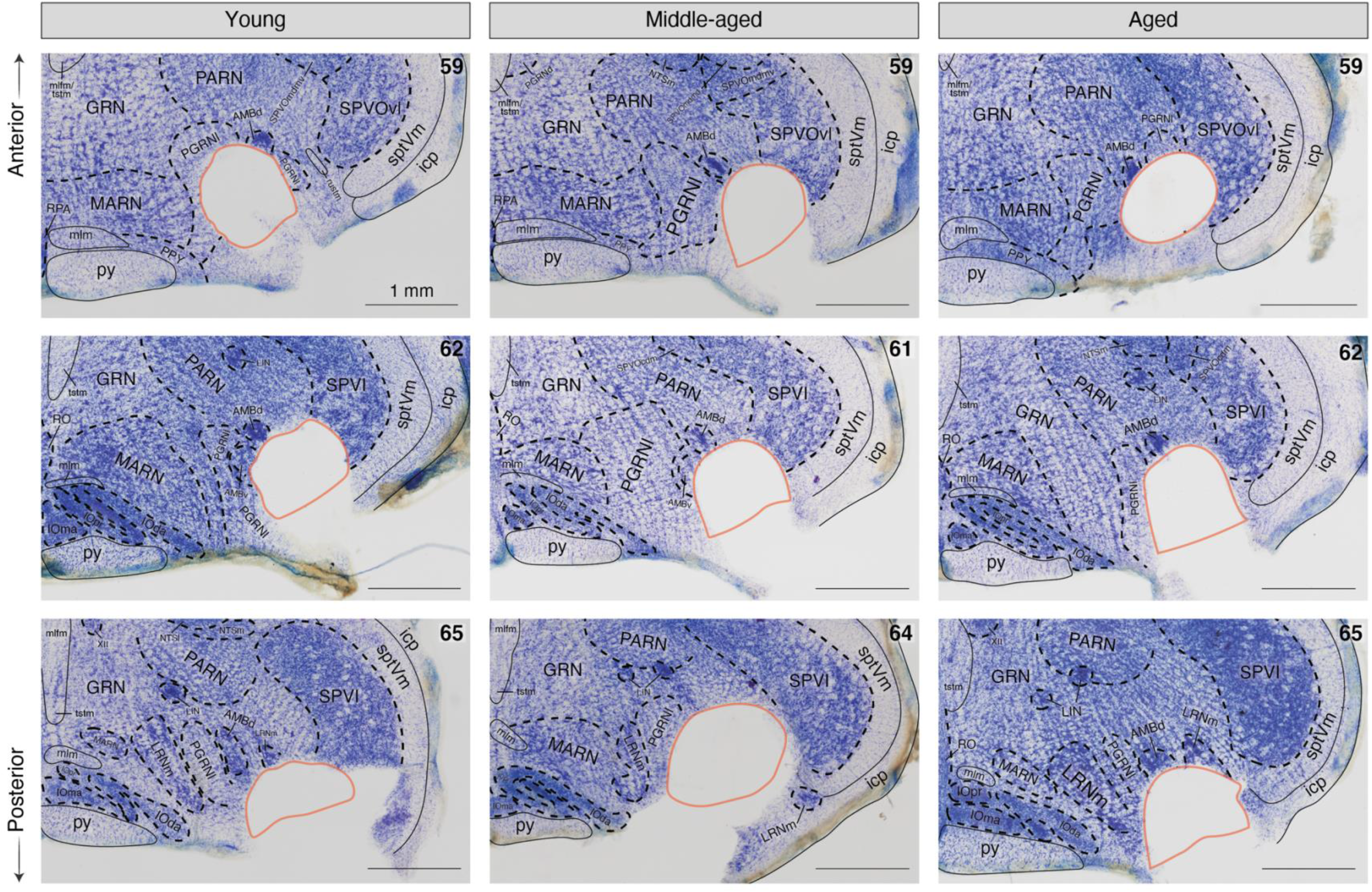
Photomicrographs displaying Nissl-based cytoarchitecture of micropunched brainstem sections. RVLM-containing area micropunched sections were stained with cresyl violet solution and representative photomicrographs from young, middle-aged, and aged groups are displayed. Gray matter region and fiber tract parcellations were contoured with dashed and solid black lines, respectively. Age group labels are displayed at the top of the figure and corresponding *BM4.0* atlas level numbers are displayed at the top right of each image. Scale bars denote one millimeter.

**Figure 10:**
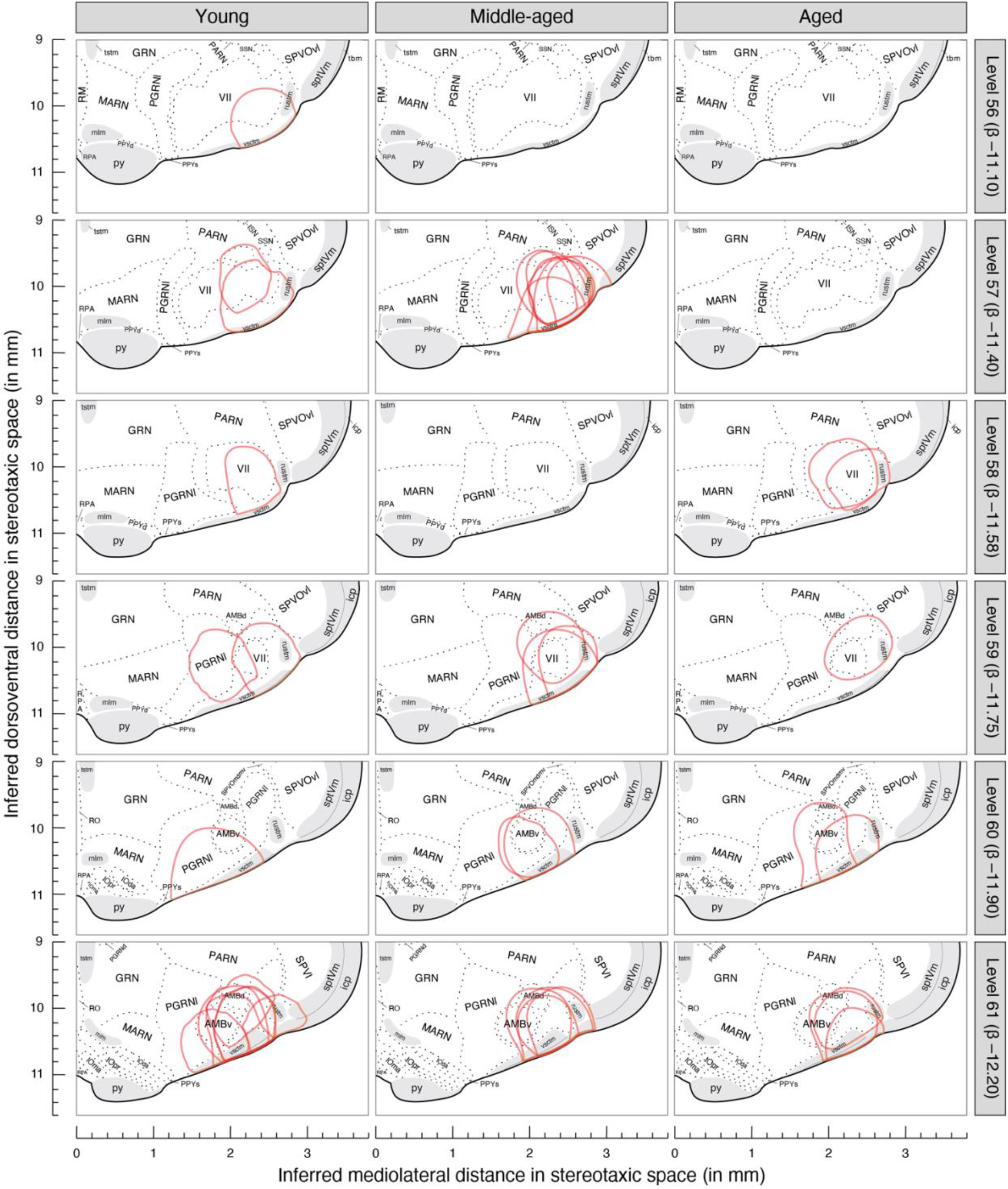
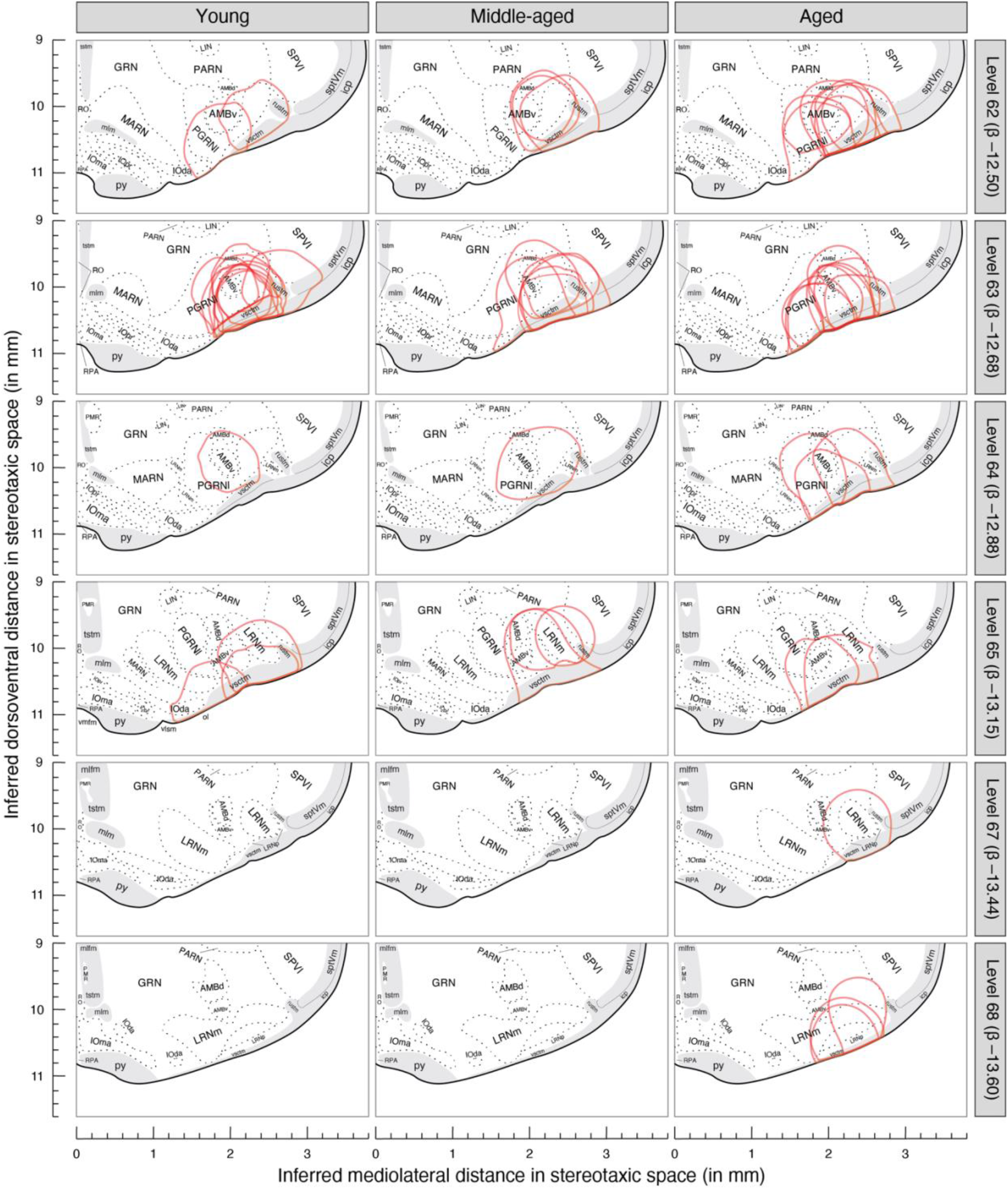
Mapping of the micropunched sites onto *Brain Maps 4.0 (BM4.0)* atlas level templates. Micropunched sites from young, middle-aged, and aged rats were mapped onto *BM4.0* atlas level templates 56–65, 67, and 68. Maps were paneled with age groups as columns and atlas levels as rows. For visualization purposes, each section’s left and right-side locations of the micropunches were mapped onto the right side of the atlas template. The outline of each micropunched site was traced in red color. Inferred mediolateral and dorsoventral distances in stereotaxic space are represented on the x- and y-axis, respectively. Age group labels are displayed at the top of the figure. Atlas level numbers are displayed at the right side of the figure and inferred Bregma distance (β) for each atlas level are reported in the parentheses (in mm). On the maps, gray matter regions are bordered with dashed lines and white matter tracts are shaded with gray background. Sample size for the Young group: 29 micropunches (9 rats), Middle-aged group: 29 micropunches (11 rats); and Aged group: 32 micropunches (8 rats).

The anterior-posterior density distribution of the sampled sites is presented as a density curve for each age group in **Figure 11** and atlas levels are represented as inferred distances from Bregma (β) on the x-axis. Tail probabilities for empirical cumulative densities are color-coded on these density curves (**Fig. 11**). For all the age groups, peak density was observed at or around atlas level 63 (−12.68 mm from β). In addition, we observed a second peak at level 57 (−11.40 mm from β) in the middle-aged group. According to *BM4.0*, the RVLM may be centered in the *paragigantocellular reticular nucleus, lateral part (>1840)* (PGRNl), and the *ambiguus nucleus ventral division (>1840)* (AMBv), corresponding to the ventrolateral area between atlas levels 60–66 (between −11.90 to −13.28 mm from β). The empirical probabilities observed on the density curves between −11.90 and −13.28 mm were 0.8, 0.6, and 0.8 for young, middle-aged, and aged groups, respectively. For direct comparison, areas and landmarks related to the RVLM in *The Rat Brain in Stereotaxic Coordinates* (*PW7*) (Paxinos and Watson, 2014) are presented below the x-axis in **Fig. 11**. Dorsoventral and mediolateral relative density distributions of the sampled sites are presented for each age group at each atlas level in **Figure 12** (blue-low, green-medium, and red-high). In young and aged rats, a relatively high sampled site density was observed at atlas level 63 (**Fig. 12** left and right panels, green to red). In middle-aged rats, a relatively high sampled site density was observed at atlas levels 57 and 63 (**Fig. 12** middle panel, green to red). In all three age groups, at level 63, the highest density area was observed in the AMBv and lateral PGRNl regions (**Fig. 12** Level 63, red).

**Figure 11:**
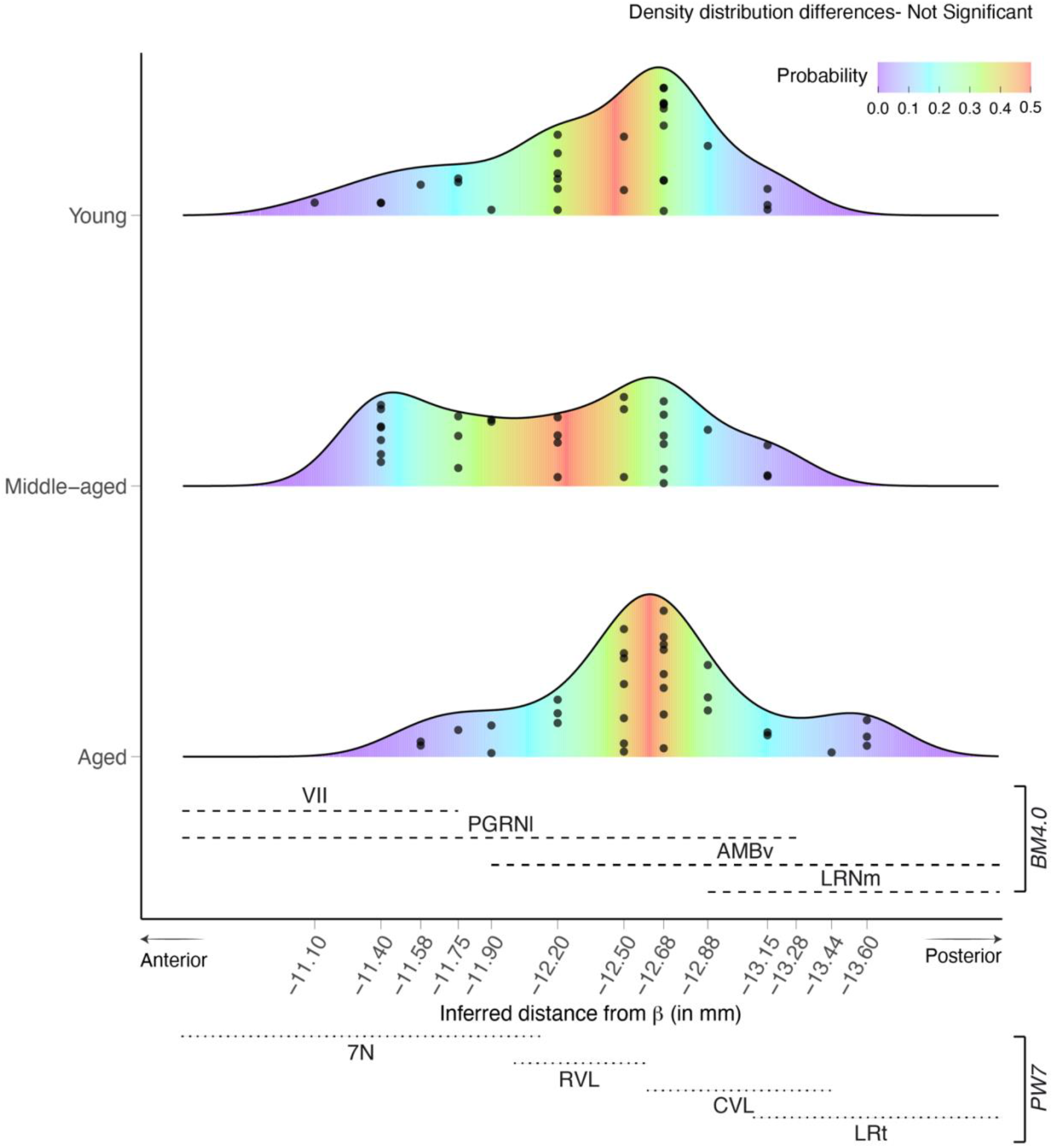
Density distribution of the micropunched sites along the anteroposterior axis. Each micropunched site’s anterior-posterior location was identified from its corresponding *BM4.0* atlas level template as an inferred distance from Bregma (β), and the anterior-posterior distribution of all the sampled sites is presented as a density curve for each age group. Areas under the curve were color-mapped with associated cumulative density tail probabilities (key-top right). In addition, each sampled site is presented as a black dot on top of the density plot. Density distribution differences between age groups are not significant at *p* < 0.05 (two sample K-S test). Reference regions from *BM4.0* are presented at the top of the *x*-axis and their anterior-posterior distribution is rendered with dashed lines. VII, Facial nucleus (>1840); PGRNl, *Paragigantocellular reticular nucleus, Lateral part (>1840)*; AMBv, *Ambiguus nucleus ventral division (>1840)*; and LRNm, *Lateral reticular nucleus, Magnocellular part (>1840)*. In addition, reference regions from the stereotaxic coordinate system of the rat brain atlas by Paxinos & Watson (*PW7*) were presented below the *x*-axis and their anterior-posterior distributions are rendered with dotted lines. 7N; facial nucleus, RVL; rostroventrolateral reticular nucleus, CVL; caudoventrolateral reticular nucleus, LRt; lateral reticular nucleus.

**Figure 12:**
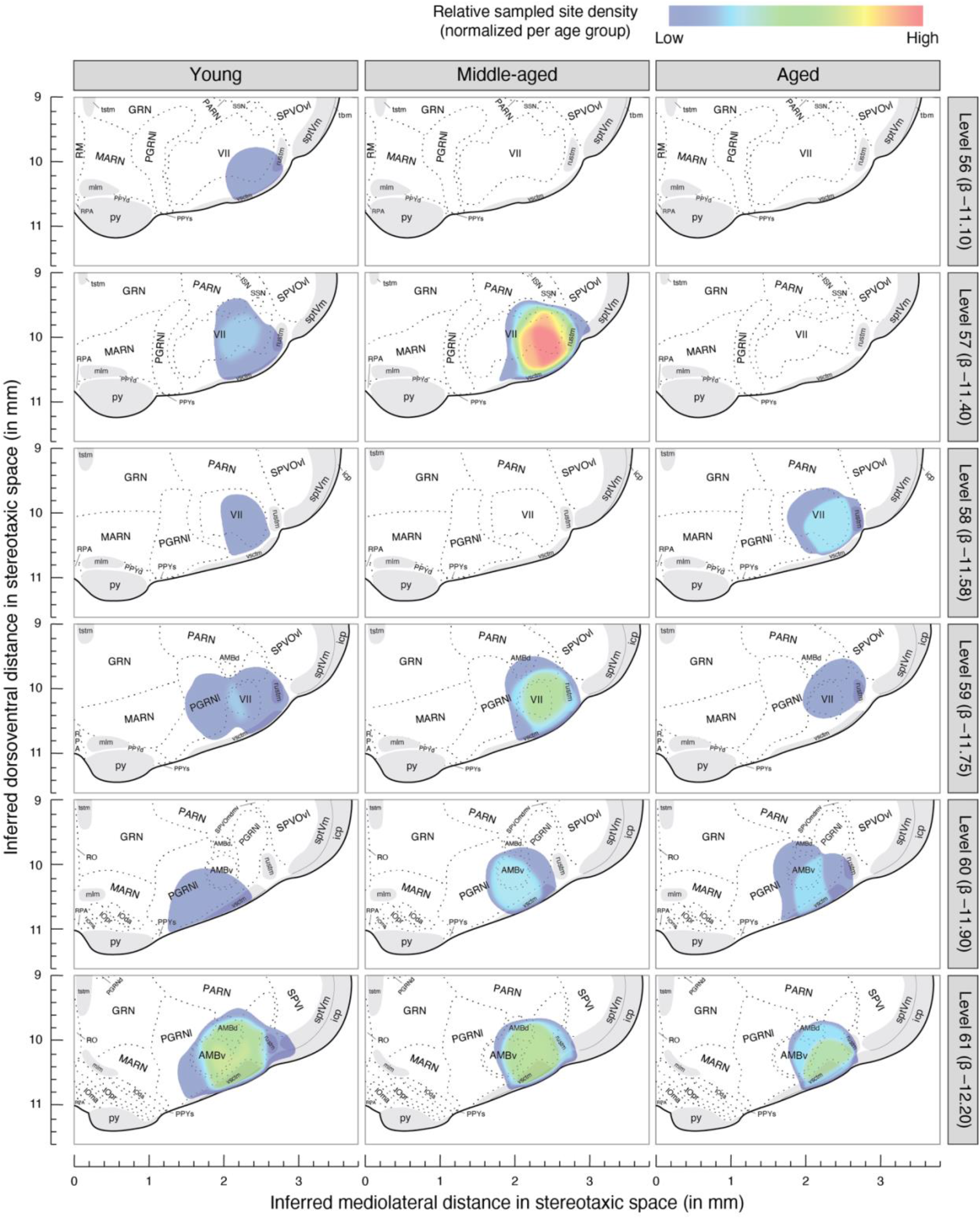
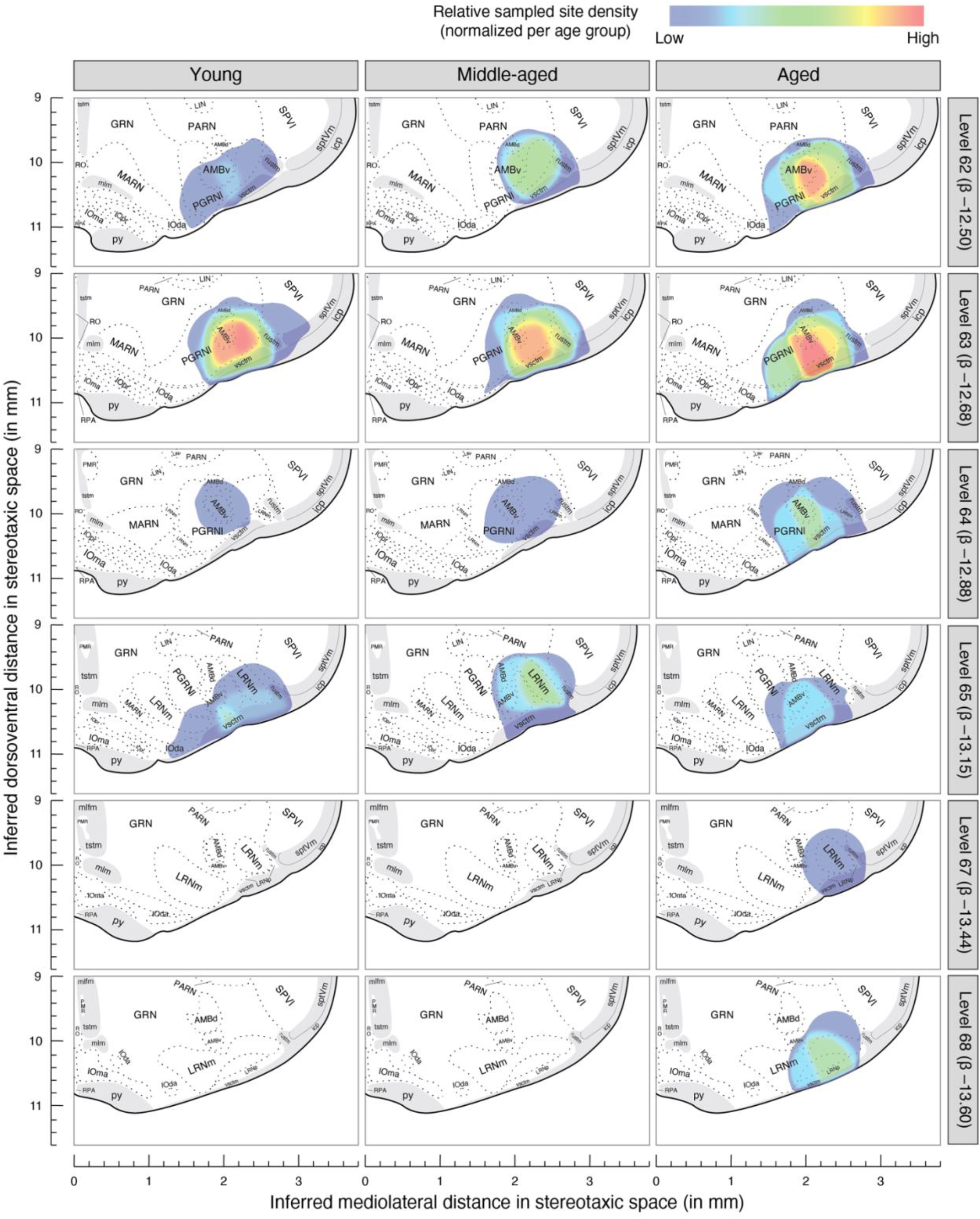
Density distributions of the micropunched sites in dorsoventral and mediolateral directions. Relative sampled site density for each atlas level was color coded and presented on their corresponding atlas level templates for each age group. At any spatial location on any atlas level template, a higher number of the sampled site tracings and their spatial overlap results in high sampled site density, and a lower number of sampled site tracings and/or their lack of spatial overlap results in low sampled site density. The color key is at the top right of the figure. Densities were normalized for each age group. Inferred mediolateral and dorsoventral distances in stereotaxic space are represented on x- and y-axis, respectively. Age group labels are displayed at the top of the figure. Atlas level numbers are displayed at the right side of the figure and inferred Bregma distance (β) for each atlas level is reported in parentheses. On the maps, gray matter regions are bordered with dashed lines and white matter tracts are shaded with gray background. Sample size for Young group: 29 sampled sites (9 rats); Middle-aged group; 29 sampled sites (11 rats); and Aged group: 32 sampled sites (8 rats).

## Discussion

In this study, we tested the hypothesis that aging is associated with altered GABA and/or glutamate neurotransmission-related gene expression in the RVLM of Fischer 344 rats. Three sets of findings support this hypothesis. First, the RVLM of middle-aged and aged rats was characterized by decreased GABA and glutamate neurotransmission-related gene expression at the RNA level compared to young rats. The observation that all of the significantly-altered genes have a downregulated expression in the RVLM of middle-aged and aged rats supports this finding. In addition, most of the remaining analyzed genes showed a downregulatory trend in middle-aged and aged RVLM. Second, among GABA and glutamate neurotransmission-related genes, the aging-associated downregulatory effect appears to be more prominent for the expression of glutamate neurotransmission-related genes in the RVLM. The observation that there were more statistically significant downregulated neurotransmission-related genes for glutamate than GABA at the RNA level in the RVLM of middle-aged and aged rats, and the downregulatory trend observed for high-affinity glutamate transporter EAAT4 (*Slc1a6*) protein expression in the middle-aged and aged RVLM, supports this finding. Several of these glutamate neurotransmission-related gene products, such as AMPA and NMDA glutamate receptor subunits, play a key role in regulating glutamatergic tone at the RVLM. Third, the effect of aging on RVLM GABA and glutamate neurotransmission-related gene expression appears to be initiated by middle-age and sustained in aged F344 rats. The absence of spatial separation between middle-aged and aged arrays in PCA analysis and the similar downregulatory pattern observed in the middle-aged and aged RVLM compared to the young RVLM supports this finding. These observations are consistent with our previous findings, where we observed an altered chemical neurotransmission-related gene expression in the aged RVLM (Balivada et al., 2017; Pawar et al., 2018), but also extends these findings by identifying an age-associated downregulation in numerous genes related to the two major amino acid neurotransmission systems (GABA and glutamate) that tonically influence RVLM activity.

A functional balance of GABAergic and glutamatergic tone in the RVLM regulates cardiovascular and sympathetic nervous system activity under basal conditions (Ito and Sved, 1997; Kenney, 2014; Kenney et al., 2011; Ross et al., 1984; Sun and Guyenet, 1985), and the current study observed an age-associated downregulation of numerous genes involved in GABA and glutamate neurotransmission in RVLM-containing tissue. Specifically, the present results indicate that of the GABA and glutamate neurotransmission-related genes analyzed, 29% and 38% were significantly downregulated in the RVLM of aged and middle-aged rats, respectively. In addition, the majority of the remaining analyzed genes displayed a chronological age-associated downregulatory trend.

All significant gene expression reductions in aged and middle-aged rats were less than two-fold in magnitude, which is generally considered to be the lower end of the qPCR analysis. Although the probability of finding small-magnitude down regulatory changes of multiple genes in the RVLM of *both* middle-aged and aged rats is low in the absence of actual differences in the gene expression, three experimental controls were completed to ensure validity of the data. First, accurate relative qPCR comparisons between different experimental groups are dependent on a stable background of reference gene expression (Vandesompele et al., 2002) and in the initial analytical procedural step the expression patterns of housekeeping genes were analyzed. Three separate housekeeping genes demonstrated consistent expression patterns across age groups and were used as reference genes. Importantly, similar age-related downregulatory trends in neurotransmission-related genes were observed when the threshold cycle (Ct) values were normalized with different combinations of the selected reference genes as well as when all three reference genes were combined (geometric mean), ruling out the possibility that age-related differences in the background level of reference genes contributed to gene expression changes between age groups. Second, the possibility was considered that the age-associated downregulation observed in the current study could be related to RVLM sampling differences among age groups; that is, the RVLM-containing area micropunched from young, middle-aged, and aged rats could be spatially different along the sampled anterior-posterior axis of the RVLM. To test this possibility, the validity of the RVLM micropunching method used in this study was analyzed for a separate set of micropunched samples obtained from a different set of young, middle-aged, and aged rat brains than those used for our gene expression analysis. The goal of this validation experiment was to use the same type of sampling method we used to obtain our gene expression data, but this time, map the sampled sites onto cytoarchitecture-based *Brain Maps 4.0 (BM4.0)* atlas templates (Swanson, 2018), following a standardized atlas-based mapping method (Khan et al., 2018), and check for their spatial uniformity. These *BM4.0* sampled site maps allowed us to identify the anterior-posterior level of each micropunched site in terms of inferred distance from Bregma. There were no significant age-associated anteroposterior distribution differences in RVLM sampling, suggesting age-associated differences in our gene expression data are not due to sampling errors associated with our RVLM micropunching method. Mapping micropunched sites on *BM4.0* atlas templates further validated the anatomical location of the sampled sites collected through the RVLM micropunching method used in the current study. Third, the SYBR^®^ green chemistry-based qPCR method used in RT^2^ profiler arrays has been reported to inflate results, yielding skewed gene expression profiles in the absence of proper controls (Alvarez and Doné, 2014). To reduce this possibility, the analytical protocol included quality control measures for rat genomic DNA contamination, PCR array reproducibility, and reverse transcription efficiency. In the present study, all of these variables were found to be consistent among young, middle-aged, and aged RVLM arrays. In addition, we independently validated selected gene (*Slc1a6*, *Slc6a13*) downregulation using TaqMan^®^-based qPCR. Collectively, the results of these analyses support the idea that the downregulatory pattern in GABA and glutamate neurotransmission-related genes observed in the middle-aged and aged RVLM represents an aging-associated effect.

Most of the significantly downregulated genes identified in the aged and middle-aged RVLM were related to glutamate neurotransmission and included genes that express proteins provisioning the glutamatergic synapse (13 out of 15 significantly altered genes were related to glutamate neurotransmission, KEGG pathway: rno04724) (Di Maio, 2021; Kanehisa and Goto, 2000). For example, AMPA and NMDA ionotropic (*Gria1*, *Grin1*, *Grin2a*, and *Grin2b*) and metabotropic (*Grm1*, *Grm5*, and *Grm7*) glutamate receptor subunits, and post-synaptic structure related genes such as *Dlg4*, *Homer1*, and *Homer2* were downregulated in the aged RVLM. Similarly, AMPA and NMDA, ionotropic (*Gria1*, *Gria2, Gria3, Gria4, Grin1*, *Grin2a*, and *Grin2b*) and metabotropic (*Grm1*, *Grm3, Grm5*, and *Grm8*) glutamate receptor subunits, and postsynaptic structure-related genes such as *Homer1* and *Homer2* were downregulated in the middle-aged RVLM. In addition, high-affinity glutamate transporter EAAT4 (*Slc1a6*) protein displayed a downregulatory trend in the RVLM of middle-aged and aged rats. These results suggest age-associated changes in the RVLM glutamatergic synapses, especially on the postsynaptic side (Balivada et al., 2017; Di Maio, 2021). Several of these glutamatergic synapse-associated ionotropic (*Gria1*, *Gria2*, *Gria3*, *Gria4*, *Grin1*, *Grin2a*, and *Grin2b*) and metabotropic glutamate receptor subunits (*Grm1*, *Grm2*, and *Grm3*) (Brailoiu et al., 2002; Llewellyn-Smith and Mueller, 2013) and postsynaptic protein (*Dlg4*) (Fyk-Kolodziej et al., 2021) are reportedly expressed in the RVLM area at the protein level and are known to be the primary responders to glutamate neurotransmitter. Thus, the data indicate that there may be a decline in the number of glutamatergic synapses in the aging RVLM, consistent with the age-associated complement-mediated synaptic pruning hypothesis proposed in our previous aging RVLM study (Balivada et al., 2017) and in other aging brain studies (Stephan et al., 2013). In addition, age-associated decreases in the density of glutamate receptors, especially NMDA receptors, have been observed in forebrain areas of F344 rats, including the cerebral cortex, hippocampus, and striatum (Segovia et al., 2001). It is therefore plausible to speculate that the decrease in the gene expression of glutamatergic synapses or receptors might attenuate glutamate neurotransmission in middle-aged and aged RVLM. However, the decrease in the expression of glutamate transporters observed in the middle-aged (*Slc1a1* and *Slc1a2*) and aged (*Slc1a6*) RVLM suggests the opposite. For example, a decrease in the expression of glutamate transporters might increase the availability of synaptic/extrasynaptic glutamate and, in turn, balance or increase glutamate neurotransmission.

Although not as robust as the effect of age on RVLM glutamate neurotransmission-related genes, age-associated downregulation of several GABA neurotransmission-related genes was identified in the present study in both aged vs. young and middle-aged vs. young RVLM. For example, metabotropic GABA_B_ receptor subunits (*Gabbr1* and *Gabbr2*), GABA membrane transporter (*Slc6a1*), and transaminase (*Abat*) were downregulated in the aged RVLM. In contrast, ionotropic GABA_A_ receptor type ɑ (*Gabra4*) and type γ (*Gabrg1* and *Gabrg2*) subunits, postsynaptic structure-related proteins (*Gphn*), and the GABA membrane transporter (*Slc6a13*) were downregulated in the middle-aged RVLM. The voltage-gated ion channel gene, *Cacna1b*, was observed to be downregulated in both middle-aged and aged RVLM. Collectively, these findings provide a substrate for altered GABAergic neurotransmission in the aged RVLM, consistent with age-associated changes in GABA neurotransmission observed in other areas of the aged brain (Rozycka and Liguz-Lecznar, 2017).

One of the limitations of the current study is that not all the gene expression changes were analyzed at the protein level. Although RNA expression may not always correlate with protein expression (Vogel and Marcotte, 2012), several previous studies have observed downregulation of neurotransmission and synapse-related protein expression in the aged brain (Ethiraj et al., 2021; Hernandez et al., 2018; Rozycka et al., 2019; Rozycka and Liguz-Lecznar, 2017; Segovia et al., 2001). However, comparisons of protein vs. mRNA levels need to be made in the middle-aged and aged rats to understand the effects of aging on RVLM GABA and glutamate neurotransmission. In our case, we observed an age-associated downregulatory trend in glutamate transporter EAAT4 protein expression. Irrespective of protein level changes, the downregulation of the neurotransmission-related genes at the RNA level in the middle-aged and aged RVLM represents a conserved feature that has been observed in different areas of the aging mammalian brain (Dillman et al., 2017; Loerch et al., 2008).

Age-associated gene expression changes of GABA and/or glutamate neurotransmission in forebrain areas have been used to identify potential mechanisms underpinning age-related alterations in learning and memory (Menard and Quirion, 2012; Mitchell and Anderson, 1998; Potier et al., 2010; Stephens et al., 2011). For example, Hernandez CM et al., (2018) identified an old age-associated decrease in metabotropic glutamate receptor gene *Grm3, Grm4* and *Grm5* RNA expression in the prelimbic cortex of F344 rats using the same RT^2^ profiler arrays that were used in the present study (Qiagen, GeneGlobe ID# PARN-152Z). In agreement with RNA expression, mGluR2/3 and mGluR5 protein expression was also decreased in the medial prefrontal cortex of aged rats (Hernandez et al., 2018). When these authors infused mGluR2/3- or mGluR5-specific glutamate receptor antagonists into the prelimbic cortex of young F344 rats, they observed an impaired working memory, suggesting a possible metabotropic glutamate receptor-based mechanism in prelimbic cortex for the working memory decline in old rats (Hernandez et al., 2018). Although the direction of age-associated molecular changes in GABA and/or glutamate neurotransmission depend on brain region, and the animal model and strain used, most of the analyzed forebrain areas were characterized with an age-associated decrease in GABA and especially glutamate neurotransmission-related gene expression (Rozycka and Liguz-Lecznar, 2017; Segovia et al., 2001). For example, in addition to the aforementioned aging study of prelimbic cortex, in recent literature, with the same GABA and glutamate neurotransmission related RT^2^ profiler arrays (Qiagen, GeneGlobe ID PARN-152Z) that were used in the current study, Hernandez AR et al (Hernandez et al., 2019) demonstrated an age-associated decrease in the expression of select GABA and glutamate neurotransmission-related genes in the CA3 area of the hippocampus in 20 month-old F344 × Brown Norway F1 Hybrid rats. Although not significant, a similar trend was reported for the dentate gyrus in their study (see Figure 3 of Hernandez AT et al., 2019). Analogous to these aging forebrain studies, the gene expression changes observed in the present study can be used to decipher changes in RVLM GABA and glutamate neurotransmission and their effects on physiological functions in aged animals.

In the current study, the pattern of decreased GABA and glutamate neurotransmission-related gene expression was also evident in the RVLM of middle-aged rats, and no significant differences were present between the RVLM of middle-aged and aged rats. Interestingly, in our previous global gene expression analysis, we did not observe neurotransmission-related gene expression differences between the middle-aged and young RVLM (Balivada et al., 2017). Greater sample size and sensitive qPCR methods used in the current study explain why the present results identified neurotransmission-related gene expression differences between middle-aged and young RVLM. Age-associated decreases in glutamate content, uptake and receptor expression have been reported in forebrain areas of middle-aged rats (Segovia et al., 2001). In addition, consistent with the idea of chronological age-associated changes in the RVLM and its effect on the sympathetic nervous system (SNS), our laboratory previously observed attenuated SNS responses to hyperthermia and hypothermia in middle-aged and aged rats compared to young rats (Helwig et al., 2006; Kenney and Fels, 2002).

Finally, two points concerning our anatomical analysis are worth noting here. First, the present results were linked to the reference atlas of Swanson (2018) for the rat brain, which is based on the brain of an adult male Sprague Dawley (SD) rat. The basic cytoarchitectonic regionalization of the SD brainstem is not expected to differ significantly from that of the Fischer F344 rat, especially since both the F344 and Sprague-Dawley lineages derive from the original Wistar lineage (Nadon, 2006). Nevertheless, any mapped assignments of the micropunched areas to the underlying cytoarchitectonically-defined brainstem regions in this study must be considered provisional until they can be reconciled in a Nissl-based atlas for the Fischer rat. At the time of this writing, only an MRI atlas of the F344 strain is available, which does not provide sufficient resolution to make comparisons of the RVLM-containing area with that of the Sprague-Dawley rat (Goerzen et al., 2020).

Second, the approach we used here to anchor micropunched regions to atlas-mapped areas of the brain allows for important linkages between gene expression changes and neural substrates implicated in not only autonomic functions but in the regulation of food intake and body weight. A few studies have reported “bulk” or single-cell transcriptomic surveys for brainstem regions implicated in these functions, including (in addition to the RVLM) the area postrema and/or the nucleus of the solitary tract (see, e.g., in rats: Liberini et al., 2016; Ramachandran et al., 2020; in mice: Dowsett et al., 2021). However, these studies have not linked their microdissected regions to precise locations in an atlas reference space. In contrast, the present study provides high spatial-resolution atlas-based mapping of RVLM-containing cytoarchitectonic regions displaying transcriptomic changes that are associated with aging and the regulation of autonomic function and energy balance. Given that caloric restriction increases maximum lifespan concomitant with decreased SNS activity in animals (Heilbronn and Ravussin, 2003), and that exercise has been shown to improve SNS function in healthy and overweight individuals in a randomized controlled trial (de Jonge et al. 2010), our work provides important new information on an SNS-regulating region where aging and alterations in energy balance are likely to produce synergistic changes in gene expression. It remains to be seen what the effects of caloric restriction would be on the gene expression profiles we measured here in association with aging, but the results of experiments in the Fischer 344 strain that show reduced age-related lesions in association with caloric restriction (Lipman et al., 1999) suggest that our observed gene expression changes in glutamate and or GABA neurotransmission would be associated with adaptive and not pathological or deleterious behavioral and autonomic function in aged animals.

In conclusion, the identification of age-associated GABA and glutamate neurotransmission changes in the RVLM area is important for understanding the effects of advanced age on central mechanisms regulating physiological function. With regards to cardiovascular and SNS regulation, our laboratory has previously explored the RVLM with electrophysiological (Kenney, 2014) and molecular approaches (Balivada et al., 2017; Pawar et al., 2017), and reported age-related alterations in GABA and glutamate neurotransmission in the RVLM (Balivada et al., 2017; Kenney, 2014). The present results extend our previous findings by identifying age-associated downregulation in GABA and glutamate neurotransmission-related gene expression in the RVLM, with a possible greater aging effect on the glutamate system. How age-related gene expression changes, with a particular focus on glutamatergic neurotransmission, affect the functionality of the aging RVLM needs to be tested in future studies using combined anatomical, cellular, and electrophysiological approaches. One key approach may involve determining potential age-related effects on specific neuron or glia types in the RVLM.

## Acknowledgments

The senior authors of this manuscript would like to acknowledge Dr. Chanran Ganta for his input in the experimental design. This study was supported by the funds awarded to Michael J. Kenney from the National Institute of Aging (NIA) (R01AG-041948) and to Arshad M. Khan from the National Institute of General Medical Sciences (NIGMS) (SC1GM127251).

## List of abbreviations

2Cb: lobule 2 of the cerebellar vermis
4V: 4th ventricle
7N: facial nucleus
VII: facial nucleus (>1840)
XII: hypoglossal nucleus (>1840)
AMBd: ambiguus nucleus dorsal division (>1840)
AMBv: ambiguus nucleus ventral division (>1840)
Aq: aqueduct
CVL: caudoventrolateral reticular nucleus
GRN: gigantocellular reticular nucleus (>1840)
icp: inferior cerebellar peduncle
icp [Brain Maps 4.0]: inferior cerebellar peduncle (Günther, 1786)
IOda: inferior olivary complex, dorsal accessory olive (>1840)
IOma: inferior olivary complex, medial accessory olive (>1840)
IOpr: inferior olivary complex, principal olive (>1840)
ISN: inferior salivatory nucleus (>1840)
LIN: linear medullary nucleus (>1840)
LRNm: lateral reticular nucleus, magnocellular part (>1840)
LRNp: lateral reticular nucleus, parvicellular part (>1840)
LRt: lateral reticular nucleus
MARN: magnocellular reticular nucleus (>1840)
mlfm: medullary medial longitudinal fascicle (Swanson, 2015)
mlm: medullary medial lemniscus (Swanson, 2015)
NTSl: nucleus of solitary tract, lateral part (>1840)
NTSm: nucleus of solitary tract, medial part (>1840)
ov: olivocerebellar tract
PARN: parvicellular reticular nucleus (>1840)
PGRNd: paragigantocellular reticular nucleus, dorsal part (>1840)
PGRNl: paragigantocellular reticular nucleus, lateral part (>1840)
PMR: paramedian reticular nucleus (>1840)
PPY: parapyramidal nucleus (Swanson, 1998)
PPYd: parapyramidal nucleus, deep part (Swanson, 1998)
PPYs: parapyramidal nucleus, superficial part (Fukuda et al.,1993)
py: pyramid (Willis, 1664)
RO: obscurus raphe nucleus (>1840)
RPA: pallidal raphe nucleus (>1840)
rustm: medullary rubrospinal tract (Swanson, 2015)
RVL: rostroventrolateral reticular nucleus
sp5: spinal trigeminal tract
sptVm: medullary segment of spinal tract of trigeminal nerve (Swanson, 2015)
SPVI: spinal nucleus of trigeminal nerve, interpolar part (>1840)
SPVOcdm: spinal nucleus of trigeminal nerve, oral part, caudal dorsomedial part (>1840)
SPVOmdmv: spinal nucleus of trigeminal nerve, medio dorsomedial part, ventral zone (>1840)
SPVOvl: spinal nucleus of trigeminal nerve, oral part, ventrolateral part (>1840)
SSN: superior salivatory nucleus (>1840)
tbm: medullary trapezoid body (Swanson, 2018)
tstm: medullary tectospinal tract (Swanson, 2018)
vsctm: medullary ventral spinocerebellar tract (Swanson, 2015)

